# Dynamic-Structure Redesign of Calmodulin Reveals Mechanistic Constraints on Ryr2 Regulation

**DOI:** 10.64898/2026.04.14.717973

**Authors:** Vladimir Bogdanov, Svetlana Tikunova, Nicolas Fadell, Robyn T. Rebbeck, Melanie L. Aprahamian, Md Nure Alam Afsar, Aleksei Chekodanov, Daniel J. Blackwell, Bjorn C. Knollmann, Razvan L. Cornea, Pete M. Kekenes-Huskey, Steffen Lindert, Christopher N. Johnson, Sandor Györke, Jonathan P. Davis

## Abstract

Calmodulin (CaM) is a highly conserved Ca^2+^ sensor that regulates hundreds of cellular targets through Ca^2+^ -dependent conformational dynamics. Despite its central role in Ca^2+^ signaling and disease, its evolutionary conservation and structural flexibility have suggested that CaM is resistant to rational redesign. Here, using the cardiac Ca^2+^ release channel Ryanodine receptor 2 (RyR2) as a model system, we tested whether incorporating conformational dynamics into computational protein design enables functional reengineering of CaM. We first applied a static structure–based redesign to increase CaM–RyR2 affinity. Although the resulting variant bound more tightly to both the RyR2 peptide and the intact channel in vitro, it distorted peptide geometry and worsened Ca^2+^ leak in cardiomyocytes ex vivo. Guided by molecular dynamics simulations, we then developed a dynamic-structure redesign strategy that preserves conformational integrity while strengthening binding. The resulting CaM variant exhibited increased RyR2 affinity and reduced pathological Ca^2+^ leak in a disease-relevant model. These findings show that improved binding affinity alone is insufficient to enhance physiological regulation and that successful CaM redesign requires preservation of conformational dynamics. More broadly, they demonstrate that integrating conformational dynamics into protein redesign can enable functionally predictive engineering of flexible regulatory protein–protein interactions.

## 3. INTRODUCTION

Calcium (Ca^2+^) signaling lies at the core of numerous physiological processes and is frequently dysregulated in disease^1,2^. Aberrant Ca^2+^ dynamics contribute to neurodegenerative disorders such as Alzheimer’s^3^, Huntington’s^4,5^, and Parkinson’s diseases^6^; disrupt excitation–contraction coupling in the heart, leading to arrhythmias^7,8^ and heart failure^9^; and play critical roles in tumor progression^10^. This broad involvement positions Ca^2+^ signaling as a central node in human disease. At the center of this network, calmodulin (CaM) functions as the preeminent Ca^2+^ sensor and signal transducer. CaM is a highly dynamic regulatory protein whose function emerges from its ability to sample and transition between a wide array of conformational states^11,12^. Together with spatiotemporal regulation of CaM gene expression^13^, this intrinsic flexibility enables CaM to regulate more than 300 distinct proteins through context-dependent structural adaptation^14^.

This functional versatility positions CaM as a compelling candidate for correcting pathological Ca^2+^ signaling^15–18^. Advances in gene therapy^19^, delivery platforms^20^, and protein engineering^21-23^ have raised the possibility that redesigned CaM variants could selectively modulate the function of specific targets and restore signaling homeostasis^18^. Because CaM–target interactions are structurally constrained yet strongly shaped by conformational dynamics, computational redesign offers a rational way to search sequence space for variants that improve target-specific regulation. Paired with experimental validation, this approach can test whether such variants preserve the dynamic behavior required for physiological function. Early pioneering computational approaches demonstrated that CaM could be redesigned with some target specificity; however, this was achieved largely by decreasing CaM’s affinity for “off” targets^24-27^. Together with CaM’s extraordinary evolutionary conservation—three genes encoding an identical protein sequence across mammals^28^— these findings highlighted the stringent constraints on functional redesign of CaM. Yet naturally occurring CaM variants in plants^29^ and engineered mammalian CaM mutants with altered Ca^2+^ sensitivity^18^ demonstrate that sequence variation can be tolerated without loss of physiological function, suggesting that the principal constraint on CaM redesign may be preservation of regulatory dynamics^30,31^ rather than strict sequence immutability. We therefore hypothesized that CaM could be *in silico* reengineered to enhance target-specific regulation if its dynamic structural constraints were explicitly incorporated into the redesign process^23, 32-34^.

To test this hypothesis, we focused on the cardiac ryanodine receptor (RyR2), a Ca^2+^ release channel whose regulation by CaM depends on domain-specific, Ca^2+^-dependent conformational transitions^35-39^ (**Fig. 1A**). Impaired CaM–RyR2 interaction destabilizes channel gating, leading to pathological diastolic Ca^2+^ leak and arrhythmia^37-39^. Because the mechanistic basis of CaM-mediated RyR2 regulation is well characterized^18,39-40^, this system provides a tractable model for testing whether redesigns that preserve dynamic integrity can translate enhanced affinity into improved physiological regulation. Here, we introduce a dynamic-structure redesign framework (**Fig. 1B**) that integrates static structural modeling with molecular dynamics simulations and functional validation. Using this approach, we demonstrate that enhanced binding affinity alone is insufficient to ensure improved physiological regulation. Instead, successful CaM reengineering requires preservation of conformational integrity within the CaM–RyR2 complex, linking structural dynamics directly to functional outcome. More broadly, our findings establish that static energy-based redesign alone may fail for flexible regulatory proteins whose activity emerges from conformational ensembles^23,32,33,41^. By explicitly incorporating dynamic constraints into the redesign process, we selectively enhanced CaM–RyR2 regulation and corrected pathological Ca^2+^ handling *ex vivo* in a disease-relevant model.

**Figure 1.**
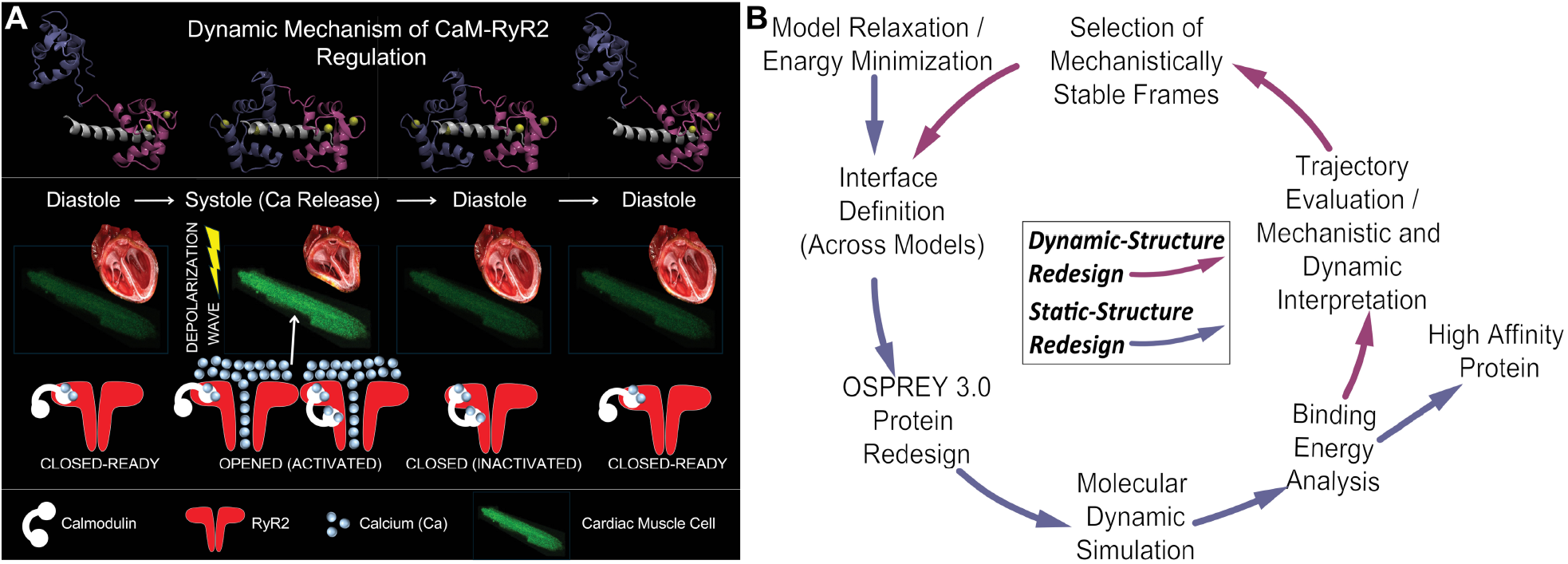
Mechanism of CaM-Mediated RyR2 Regulation and Dynamic-Structure Redesign. (A) *Dynamic model of CaM–RyR2 regulation across the cardiac cycle. Top:* Modeled conformational transitions of the CaM–RyR2 complex depict rearrangements of the N-domain (blue) and C-domain (magenta) during systolic activation and diastolic recovery, with Ca^2+^ ions shown in yellow and RyR2 peptide in white. *Bottom:* Schematic representation of Ca^2+^-dependent RyR2 gating in cardiomyocytes. During diastole, the N-domain of CaM dissociates from RyR2, contributing to the channel’s *closed-ready* state. Depolarization triggers Ca^2+^ release through RyR2 (*opened/activated* state), leading to Ca^2+^-dependent saturation of the N-domain of CaM, which promotes its rebinding to RyR2, thereby stabilizing the *closed/inactivated* state essential for limiting Ca^2+^ leak. As Ca^2+^ is re-sequestered into the sarcoplasmic reticulum, CaM dissociates from RyR2, restoring the *closed-ready* state, in preparation for the next excitation cycle. (B) *Schematic of the Dynamic-Structure Protein Redesign framework*. The schematic illustrates a feedback-loop workflow integrating static (blue) and dynamic (magenta) redesign cycles. The process begins with model relaxation and energy minimization, followed by interface definition across multiple models and OSPREY 3.0–based protein redesign. Each redesigned complex undergoes molecular-dynamics simulation and energetic evaluation using MM/PBSA analysis. Insights from molecular dynamics trajectories evaluation and mechanistic interpretation of CaM–RyR2 structural dynamics guide the selection of dynamically persistent, mechanistically relevant frames, which are then used to redefine the interface for the next redesign cycle. Iterative convergence of these static and dynamic rounds yields higher-affinity, dynamically stabilized CaM variants that incorporate the intrinsic dynamic properties of the CaM–RyR2 complex, enabling the redesign to translate into measurable physiological improvement.

## RESULTS

### tatic-Structure Redesign (in silico)

Based on our current model of CaM–RyR2 regulation, in which the N-terminal regulates activity and the C-terminal anchors CaM to RyR2^18,42^ (**Fig. 1A)**, we hypothesized that strengthening the affinity of CaM’s N-domain for RyR2 would minimize pathological Ca^2+^ leakage. To test this hypothesis, we employed an atomic structure of Ca^2+^CaM-RyR2 peptide complex^43^ (PDB ID: 6Y4O). Analysis of the CaM–RyR2 interface identified 24 N-domain residues within 7Å of the peptide (**Fig. 2A**). Most of these contacts were polar or hydrophobic, distributed across the N-lobe. Energy calculations using protein redesign software^44^ (OSPREY 3.0) predicted only a small subset of substitutions to lower the complex’s energetic profile (**Fig. 2A**). Given that CaM binds hundreds of proteins with highly diverse binding motifs, this result aligns with the view that CaM’s sequence is nearly evolutionarily optimized^24^. Strikingly, only two residues—S38 and Q41—were consistently identified as suboptimal, with OSPREY predicting that substitution with glutamic acid would be more favorable (**Fig. 2A**).

**Figure 2.**
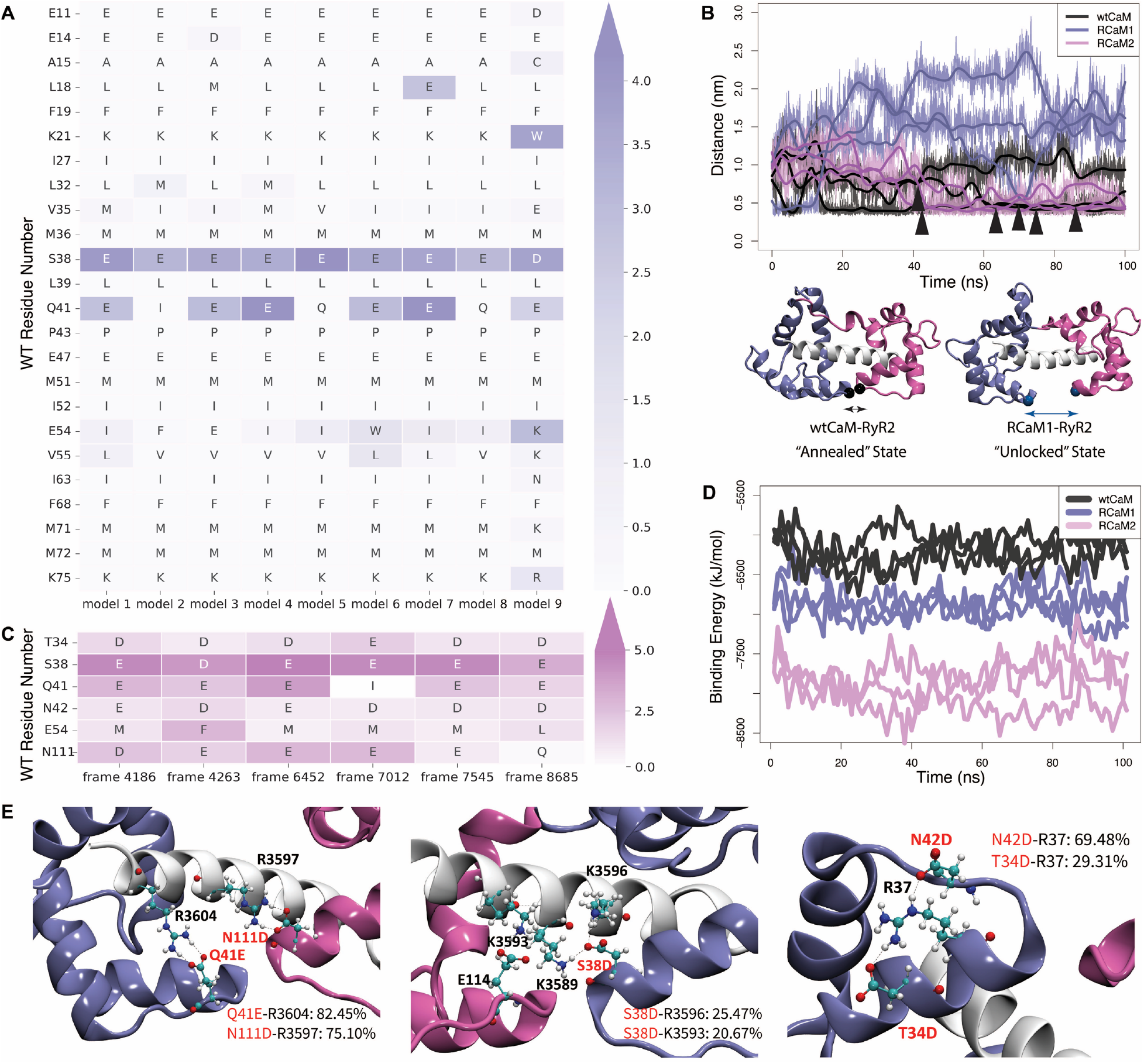
*In Silico* CaM Redesign Strategy. (A) *Static-Structure Redesign*. OSPREY 3.0 computational redesign of the CaM N-domain–RyR2 interface identified residues within 7 Å of the RyR2 peptide across nine models. Two positions—S38 and Q41—were consistently predicted as suboptimal, with both Glu and Asp substitutions predicted to improve binding energetics; only the top-ranked variants are shown. Color bar indicates ΔK* energy scores (higher = more favorable). (B) *RyR2-CaM complex dynamics*. Time-dependent distance between the N-domain residue Cα (S40) and the C-domain residue Cα (E113) demonstrates that wtCaM favors the annealed state (black, low distance), whereas RCaM1 adopts an unlocked conformation (dark blue, high distance). Representative structures below illustrate peptide bending in the unlocked RCaM1 complex versus the straight peptide in wtCaM. The RyR2 peptide is shown in white; CaM N- and C-domains of CaM are highlighted in blue and magenta, respectively. Black and dark blue spheres indicate Cα atoms of residues 40 and 113 in RCaM1 and wtCaM. Ca^2+^ ions were removed for clarity. Black arrows indicate frames corresponding to annealed states in wtCaM–RyR2 trajectories that were selected for the second-round redesign analysis. (C) *Dynamic-Structure Redesign*. From molecular dynamics trajectories of wtCaM–RyR2, six frames representing the annealed state (close N- and C-domain coupling) were selected for a second redesign round focused on residues T34, S38, Q41, N42, E54, and N111. Acidic substitutions (Glu or Asp) at T34, S38, Q41, N42, and N111 were predicted to reinforce N-domain contacts with the RyR2 peptide, whereas E54 was replaced with Met to reduce competition within the bonding network; only the top-ranked variants are shown. (D) *Binding energy analysis*. MM/PBSA energy trajectories for wtCaM, RCaM1, and RCaM2 show that both redesigned CaMs maintain lower total energy (higher affinity) relative to wtCaM, with RCaM2 exhibiting the greatest stability. Calculations were performed every 1 ns along molecular dynamics trajectories. (E) *Representative bonding networks within the RCaM2–RyR2 complex. Left:* Q41E and N111D form stable hydrogen bonds with R3604 and R3597. *Middle:* S38D interacts with K3596 and K3593. *Right:* N42D and T34D interact intramolecularly with R37, contributing indirectly to complex stabilization. Percentages indicate bond occupancy across molecular dynamics trajectories.

Although these mutations were predicted to minimize the energetic profile of the CaM–RyR2 complex, the redesign was based on static snapshots^31,45^, whereas CaM is highly dynamic^11,12,18,46^. Therefore, both the redesigned CaM (RCaM1) and wtCaM complexes were further minimized, and molecular dynamics simulations were performed. Interestingly, in most wtCaM trajectories the N- and C-domains rapidly converged and then remained in close proximity (**Fig. 2B: bottom**), a conformation we termed the “annealed” state. This motion was quantified (**Fig. 2B: top**) by tracking the distance between the Cα of residue 40 in the N-domain and the Cα of residue 113 in the C-domain for both RCaM1 and wtCaM. By contrast, RCaM1’s N- and C-domains moved apart, producing a dramatic “unlocked” state (**Fig. 2B**). Notably, this dynamic separation of RCaM1 domains markedly bent the RyR2 peptide at residue S3592, whereas the annealed state of wtCaM preserved peptide structural integrity (**Fig. 2B, Supplementary Fig. S1, Supplementary Movie SM1**). To assess whether these dramatic conformational changes and peptide bending in the RCaM1 complex influenced affinity, each trajectory from both wtCaM and RCaM1 simulations was analyzed using MM/PBSA^47^. Surprisingly, MM/PBSA revealed that despite its greater flexibility and peptide bending, the RCaM1 complex consistently remained in a lower-energy, higher-affinity state than wtCaM (**Fig. 2D**). As expected, most residue-level MM/PBSA energy differences mapped to the S38E and Q41E substitutions, with smaller contributions from RyR2 peptide residues (**Supplementary Fig. S2**). Thus, our *in silico static-structure redesign* (**Fig. 1B**) produced RCaM1, a CaM variant that not only reshaped complex dynamics—frequently sampling an unlocked state and bending the RyR2 peptide—but also paradoxically increased binding affinity relative to wtCaM.

### Dynamic-Structure Redesign (in silico)

Because the *static-structure redesign* was based on static, energy-minimized snapshots, it could not account for conformational preferences emerging from molecular dynamics^33,48^. Based on our observation that wtCaM dynamically favors an annealed conformation with a straighter RyR2 peptide (**Fig. 2B, Supplementary Fig. S1**), we rationalized that maintaining peptide integrity—rather than merely maximizing static binding energy—could be essential for proper CaM–RyR2 regulation. We therefore iterated our redesign strategy (**Fig. 1B**), focusing specifically on reinforcing the annealed state, which is absent from statically resolved or predicted RyR2–CaM structures. From molecular dynamics trajectories, we selected six representative annealed-state frames of wtCaM (**Fig. 2B: arrows**) and identified amino acids within 7Å of the RyR2 peptide. Two residues, T34 and N42—previously not considered part of the RyR2 contact surface (**Supplementary Fig. S3A, B: right**)—were now within 7 Å in the annealed wtCaM complex (**Supplementary Fig. S3A, B: left)**. Additionally, we found that residue N111 in the C-domain of CaM might be important for stabilizing annealing between the N- and C-domains (**Supplementary Fig. S3D, E**). Furthermore, E54, ranked by OSPREY in the first round of redesign to be the 3^rd^ least favorable native amino acid (**Fig. 2A**) also remained within 7Å of RyR2 in the annealed state (**Supplementary Fig. S2C**). Consequently, we initiated another round of redesign using annealed state structures of the wtCaM complex, adding an iterative cycle to our overall redesign framework (**Fig. 1B**).

To save computational time and test our rationalization, the second round of the redesign focused exclusively on residues 34, 38, 41, 42, 54 and 111 (**Fig. 2C**). Similar to the first round’s calculations, residues 41 and 38 still showed a high preference for glutamic and aspartic acids. Similarly, residues 34, 42 and 111 also indicated a preference for acidic amino acids with marginal differences in Δ*K*^*\**^*scores* between Asp and Glu (data not shown). Consistent with our first round of redesign, where we produced RCaM1, the acidic residue at position 54 was suggested to be more hydrophobically favorable (**Fig. 2A, C**). Considering OSPREY calculations and hypothesizing that shorter side chains might enhance the hydrogen bonding network between CaM and RyR2, RCaM2 was assigned the following mutations: T34D, S38D, Q41E, N42D, E54M and N111D.

Calculations on the molecular dynamics trajectories of the RCaM2-RyR2 complex, demonstrated that the overall lifetime of the bonding network increased significantly for RCaM2, surpassing those observed for both the RCaM1 and wtCaM complexes (**Supplementary Fig. S4**). Specifically, in the RCaM2 complex, the major contribution to the N-domain bonding network with RyR2 again came from S38D and Q41E, lasting ∼50% and ∼80% of the simulation time (**Fig. 2E, Supplementary Fig. S4: clusters a, b**). Interestingly, in RCaM1 E54 competed with Q41E for interaction with R3604, whereas in RCaM2, replacing M54 allowed much more stable bonding between Q41E-R3604 and N111D-R3597 (**Fig. 2E, Supplementary Fig. S4: clusters b, c**). Surprisingly, both T34D and N42D showed little direct contribution to the CaM-RyR2 bonding network. Instead, they interacted with R37 within RCaM2, yet still contributing favorably to the overall binding energy between RCaM2 and the RyR2 peptide (**Fig. 2E, Supplementary Fig. S4)**. Furthermore, the RyR2-RCaM2 complex predominantly remained in the annealed state and the RyR2 peptide remained straight and rigid during the molecular dynamics simulations (**Fig. 2B, Supplementary Fig. S1**). Consistent with OSPREY predictions, MM/PBSA analysis of RCaM2 molecular dynamics trajectories indicated a lower binding energy compared to both wtCaM and RCaM1 over time for the RyR2 peptide (**Fig. 2D, Supplementary Fig. S5**). Thus, using our *dynamic-structure redesign* strategy (**Fig. 1B**) we engineered RCaM2 with the N-domain “locked” in the annealed state, which resulted in a predicted higher affinity to RyR2 than both wtCaM and RCaM1. Overall, we computationally redesigned two CaM variants with distinct dynamic states (RCaM1: unlocked state; RCaM2: “locked” in annealed state through a reinforced bonding network) both of which were predicted to have an improved affinity to the RyR2 peptide. Subsequently, we introduced the mutations into a CaM expression plasmid and recombinantly expressed and purified each CaM construct to experimentally test these *in silico* predictions.

### Affinity Validation using the RyR2 peptide (in vitro)

To directly test our computational predictions, we examined *in vitro* whether the redesigned CaMs bind more tightly to the RyR2 CaM-binding peptide. Specifically, we followed the intrinsic tryptophan (Trp) fluorescence of the CaM-binding peptide of RyR2, which increases upon CaM binding^18^ **(Fig 3A;**). In the absence of Ca^2+^, none of the CaMs influenced the Trp fluorescence of the RyR2 CaM binding peptide (data not shown), consistent with the very low apo-state affinity of this isolated peptide^18^. However, in the presence of saturating Ca^2+^ (pCa 4.0), Trp fluorescence increased with similar apparent Kd values of ∼120 nM for all three CaMs. Since the concentration of CaM needed to measure the fluorescence is substantially above the expected Kd (∼1nM)^49^ (**Supplementary Fig. S6**) it was not surprising the apparent Kds were similar at saturated Ca^2+^. To circumvent this technical issue and considering *in vivo* Ca^2+^ levels in the cell also occur in the nM to low uM region (not saturating), we performed the RyR2 peptide steady-state binding studies also at a pCa of 6.6 (∼250 nM). Under these experimental conditions, wtCaM displays an apparent Kd for the RyR2 peptide of ∼454 nM, ∼4-fold weaker than that observed in saturating Ca^2+^ (**Fig. 3A, Supplementary Fig. S6**). Compared to wtCaM, both RCaM1 and RCaM2 display ∼3- and 4-fold higher apparent sensitivities for the RyR2 peptide, respectively (**Fig 3A**). Interestingly, decreasing the concentration of free Ca^2+^ had essentially no effect on the apparent sensitivity of RCaM2 for the RyR2 peptide (**Fig. 3A, Supplementary Fig. S6**), indicating it possesses very high affinity, even under sub-saturating Ca^2+^ conditions.

**Figure 3.**
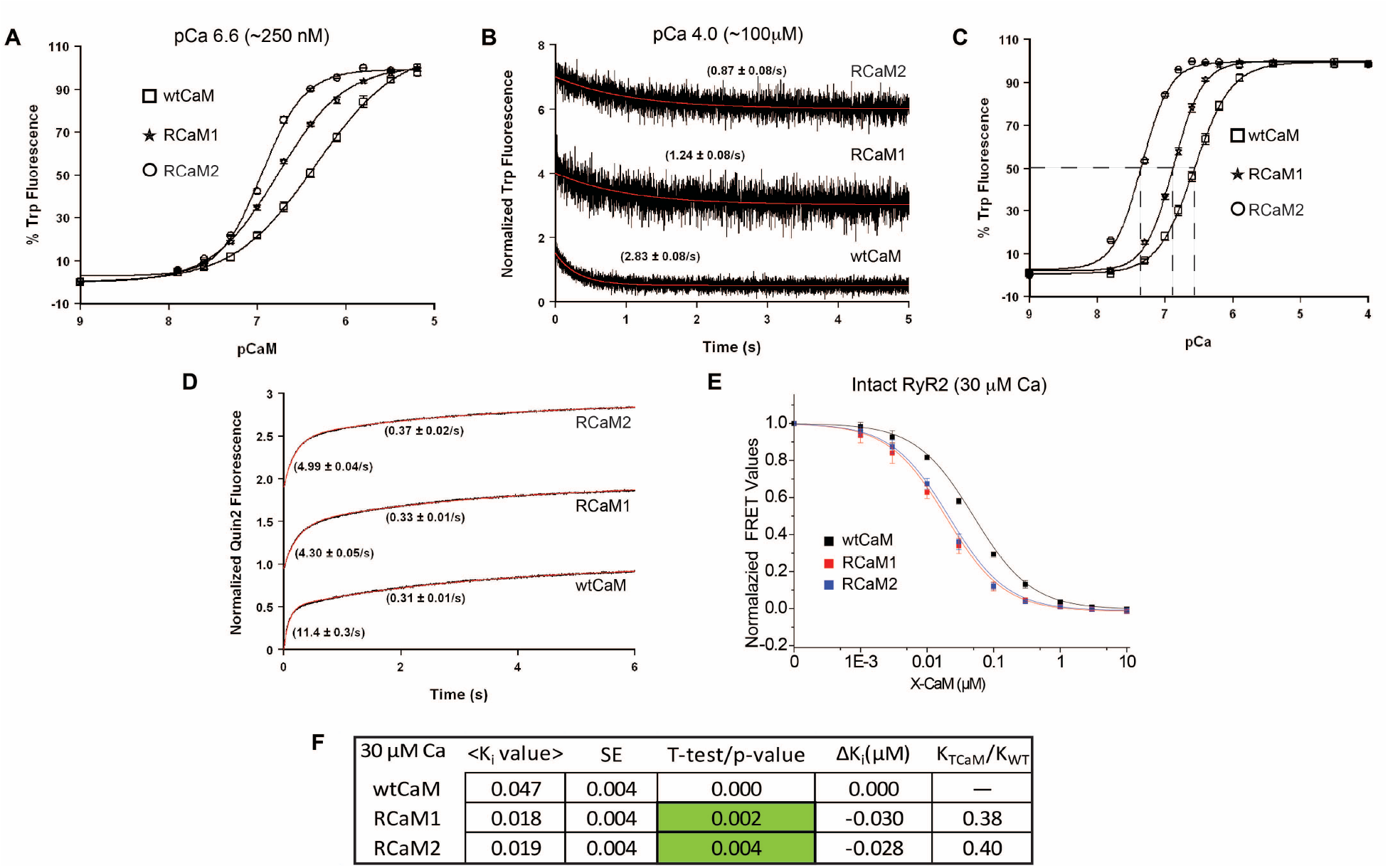
*In Vitro* Validation of the Redesign Strategy. (A) *Affinity for the RyR2 peptide at low Ca*^*2+*^. Steady-state binding of CaM variants to the RyR2 CaM-binding peptide, monitored by intrinsic tryptophan (Trp) fluorescence at sub-saturating Ca^2+^ (pCa 6.6, ∼250 nM). (B) *Dissociation rate of RyR2 peptide from CaM variants*. Peptide-competition kinetic-exchange assay monitoring Trp fluorescence decay following rapid mixing of preformed RyR2 peptide–CaM complexes with excess spectroscopically silent competitor peptide (NaV1.5). (C) *Ca*^*2+*^ *sensitivity of CaM variants*. Apparent Ca^2+^ sensitivity determined from changes in intrinsic Trp fluorescence as Ca^2+^-CaM binds the RyR2 peptide. (D) *Ca*^*2+*^ *dissociation kinetics from CaM bound to RyR2*. Stopped-flow analysis of Ca^2+^ dissociation from Ca^2+^-saturated CaM–RyR2 complexes. Biphasic Ca^2+^ release measured with the fluorescent chelator Quin-2, which resolves both the N-domain (fast) and C-domain (slow) release phases. (E-F) *Affinity for intact RyR2*. FRET-based competition assay measuring binding of CaM variants to intact Ca^2+^-saturated RyR2 channels. Table (F) summarizes of apparent K_i_ values, standard errors, and derived parameters from panel E.

Complementary to the steady-state binding assay, we employed a peptide competition kinetic exchange assay to assess the dissociation rate of the CaMs from the RyR2 peptide at saturating Ca^2+^. Specifically, we rapidly mixed Ca^2+^ saturated CaM variants bound to the RyR2 peptide with a high concentration of a non-fluorescent CaM-binding peptide (corresponding to human NaV 1.5) and measured fluorescence decay as the RyR2 peptide was displaced. This assay revealed that both RCaM1 and RCaM2 dissociated from the RyR2 peptide 2- and 3-fold more slowly than wtCaM, respectively (**Fig 3B**). The slower dissociation rates demonstrate that a reduced off-rate is the principal mechanism underlying the higher peptide affinity of the redesigned CaMs.

Additionally, since Ca^2+^ binding and RyR2 peptide binding are coupled, we also measured the Ca^2+^ sensitivity for the re-engineered CaMs in the presence of the RyR2 peptide. In steady-state Trp fluorescence assays, both RCaM1 and RCaM2 displayed ∼2- and 6-fold increases in apparent Ca^2+^ sensitivity, respectively, compared to wtCaM in the presence of the RyR2 CaM-binding peptide (**Fig. 3C**). Stopped-flow experiments demonstrated biphasic Ca^2+^ dissociation kinetics in the presence of the RyR2 peptide, corresponding to the fast (N-domain) and slow (C-domain) Ca^2+^ release phases^50^. Notably, the fast phase (N-domain) was nearly 2-fold slower for RCaM1 (∼4.3 s^−1^) and RCaM2 (∼5.0 s^−1^) compared to wtCaM (∼11.4 s^−1^) (**Fig. 3D**), indicating that the N-domain stabilization directly contributes to the reduced off-rate observed in the peptide competition kinetic exchange assay. Consistent with the anchoring role of the C-domain^51^, the slow phase remained comparable across all variants (∼0.33 s^−1^).

Together, these results demonstrate that the redesigned CaMs achieve higher affinity primarily by slowing RyR2 peptide dissociation through N-domain stabilization. This mechanistic insight is consistent with our *in silico* predictions, in which the N-domains of both variants—particularly RCaM2—formed longer-lived bonding networks with RyR2 (**Supplementary Fig. S4**).

### Affinity Validation with Intact RyR2 (in vitro)

Considering the substantial conformational changes of CaM during RyR2 activation and deactivation, and to ensure that our *in silico* predicted RCaMs engage intact RyR2, we employed a FRET-based competition assay to determine the apparent affinity of the CaMs for intact Ca^2+^ -saturated RyR2^40^ in SR vesicles. In this assay, FRET occurs between a fluorescent CaM (AF568-T34C-CaM, acceptor) bound to RyR2 near fluorescent FKBP12.6 (AF488-T6-FKBP12.6, donor); as increasing concentrations of unlabeled CaM (wtCaM, RCaM1, or RCaM2) displace the fluorescent CaM probe, FRET decreases. As shown in **Fig. 3E-D**, wtCaM displaced the fluorescent CaM probe with an apparent Kd of ∼47 nM. Consistent with *in silico* predictions and peptide biochemistry, both RCaM1 and RCaM2 bound intact RyR2 with ∼2.5-fold higher affinity than wtCaM in the presence of 30 µM Ca^2+^. Thus, both redesigned CaMs exhibit higher affinity not only for the isolated RyR2 peptide but also for the native RyR2 channel complex.

### Validation of CaM-RyR2 Regulation (ex vivo)

According to our current model of CaM-RyR2 regulation^18,42^, Ca^2+^ binding to the N-domain of CaM promotes its interaction with the RyR2 CaM-binding peptide, thereby stabilizing channel closure^39^ and biasing RyR2 towards a refractory, non-leaking state following Ca^2+^ release (**Fig. 1A, Fig. 4A**)^52,53^. Because the *in vitro* experiments relied on either isolated peptide fragments or intact RyR2 channels studied outside their native cellular environment, they cannot fully capture the structural and regulatory context that governs RyR2 function and its dynamic regulation under physiological conditions. In particular, the unlocked dynamics of RCaM1 induce bending of the RyR2 CaM-binding peptide (**Fig. 2B, Supplementary Fig. S1**), a structural perturbation that may alter the kinetics of channel reopening and thereby influence refractory behavior. We therefore next examined whether these re-engineered CaMs alter RyR2 regulation under physiological conditions *ex vivo*.

**Figure 4:**
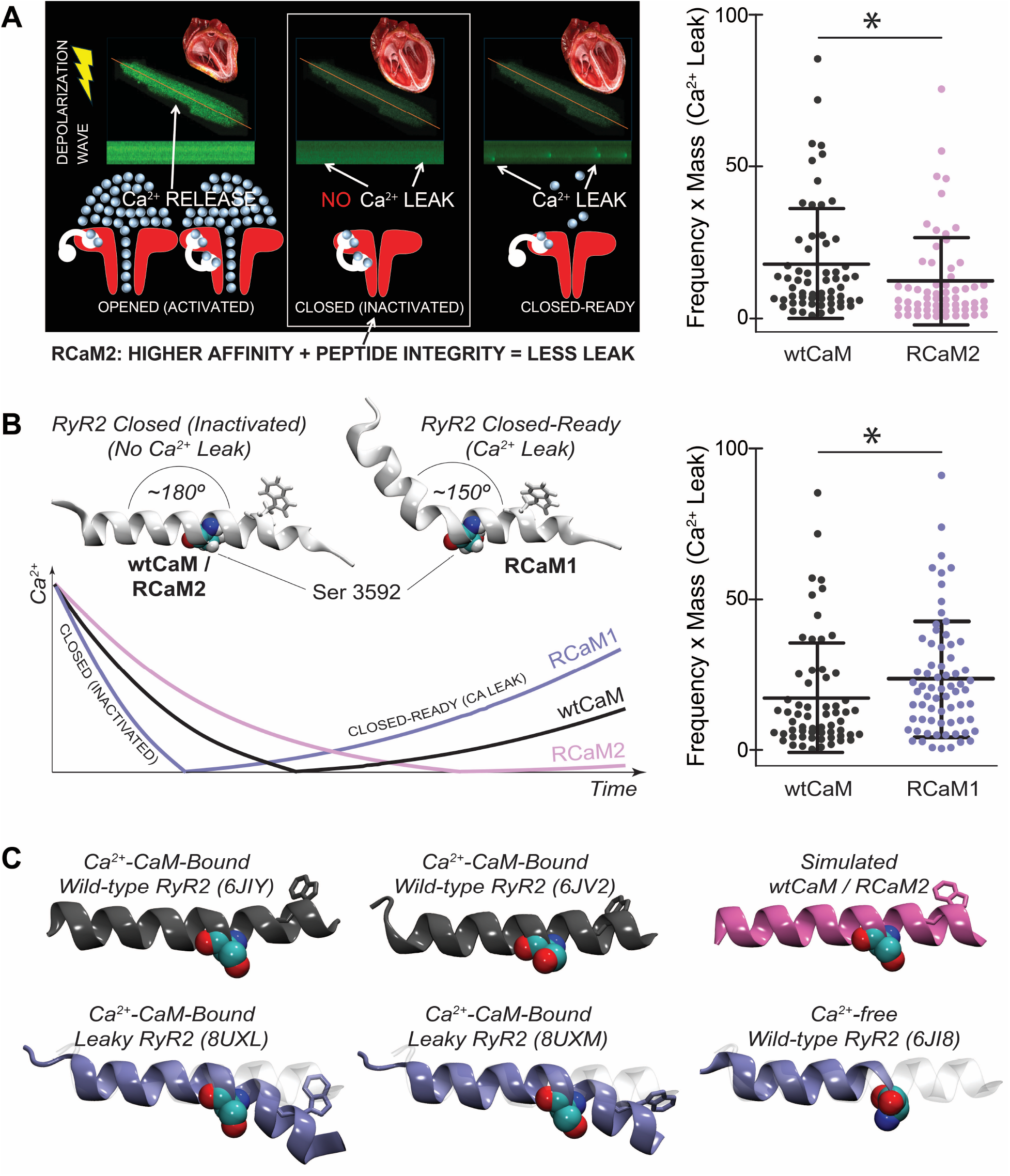
*Ex vivo* Functional Validation of the Redesign Strategy and Mechanistic Basis into CaM– RyR2 Regulation. (A) *RCaM2 suppresses pathological Ca*^*2+*^ *leak. Left:* schematic illustrating the relationship between CaM binding, RyR2 functional states, and Ca^2+^ release in the leaky RyR2 model. During diastole, spontaneous Ca^2+^ release events (leak) reflect defective RyR2 inactivation. In permeabilized cardiomyocytes, RCaM2 exhibits enhanced RyR2 association and is associated with reduced Ca^2+^ leak frequency. *Right:* quantification of spontaneous Ca^2+^ release events (frequency × mass) in permeabilized S2814D ventricular myocytes incubated with wtCaM or RCaM2 (n = 78 cells, mean ± SD, p < 0.05). (B) *Model of CaM-mediated RyR2 regulation. Left:* schematic illustrating how CaM variants influence the geometry of the RyR2 CaM-binding peptide near residue S3592. wtCaM and RCaM2 are associated with a straighter peptide orientation (∼180°), whereas RCaM1 is associated with a bent configuration (∼150°) observed in leaky states. *Bottom left:* Conceptual schematic showing how CaM variants shift RyR2 behavior along the inactivated → closed-ready (leak-prone) continuum. RCaM2 stabilizes the inactivated state, wtCaM shows intermediate behavior, and RCaM1 accelerates transition toward leak-prone conformations. Curves represent qualitative trends from simulations and ex vivo measurements, not fitted experimental data. *Right:* quantification of Ca^2+^ leak (frequency × mass) in permeabilized S2814D ventricular myocytes incubated with wtCaM or RCaM1 (n = 78 cells, mean ± SD; p < 0.05). (C) *Structural context for peptide bending near S3592*. Cryo-EM structures of Ca^2+^–CaM–bound wild-type RyR2 in the closed state (PDB IDs: 6JIY, 6JV2; gray) show a straight CaM-binding helix adjacent to S3592, consistent with the geometry observed in simulations of the wtCaM/RCaM2 complexes (magenta). In contrast, Ca^2+^–CaM–bound leaky RyR2 variants (R420W/PKA-phosphorylated forms; PDB IDs: 8UXL, 8UXM; blue) exhibit a bent helix near S3592 accompanied by a semi-transparent continuation of the CaM-binding segment. The Ca^2+^-free RyR2–apoCaM structure (PDB ID: 6JI8; blue) similarly shows an unresolved region adjacent to S3592, represented as a semi-transparent segment. W3587 is shown in licorice representation, and S3592 is displayed as space-filling spheres.

To test whether the enhanced affinity and N-domain stabilization of the re-engineered CaMs influence RyR2 regulation by prolonging its non-leaking interval, we used a mouse model carrying the RyR2-S2814D mutation, which is known to dysregulate RyR2 and promote pathological Ca^2+^ leak^54^. Saponin-permeabilized ventricular myocytes isolated from S2814D mice were incubated with wtCaM, RCaM1, or RCaM2 prior to Ca^2+^ leak assessment^18^. Surprisingly, RCaM1—predicted by both computational and biochemical assays to have improved affinity for both the RyR2 peptide and the intact channel, yet exhibiting altered dynamics that bend the peptide—increased Ca^2+^ leak compared to wtCaM (**Fig. 4B: right, Supplementary Fig. S7**). This finding suggests that increased affinity and N-domain stabilization alone are insufficient to improve RyR2 regulation. In contrast, RCaM2—combining high RyR2 affinity with dynamic features resembling wtCaM, reinforced by a stronger bonding network—significantly reduced Ca^2+^ leak in isolated cardiomyocytes (**Fig. 4A: right, Supplementary Fig. S7**). Consistent effects were observed in permeabilized ventricular myocytes expressing wild-type RyR2, where RCaM1 increased and RCaM2 decreased spontaneous Ca^2+^ spark activity (**Supplementary Fig. S8**).

Together, these findings demonstrate that effective CaM-mediated regulation of RyR2 emerges from a precise balance of affinity, domain stabilization, and dynamic integrity. Importantly, they show that even a highly conserved protein such as CaM can be computationally redesigned to restore physiological function at the cellular level in a disease-relevant model. This provides a conceptual framework for extending protein engineering strategies to other CaM-regulated channels and signaling pathways.

## DISCUSSION

Across mammals, CaM is encoded by three separate genes yet retains an identical amino acid sequence^28^, reflecting extraordinary evolutionary conservation. Together with its Ca^2+^-dependent structural adaptability and versatility in regulating more than 300 targets^14,15^, this conservation has underscored how challenging it is to redesign CaM while preserving physiological function^24^. Here, using the CaM–RyR2 complex as a model system, we show that CaM can be rationally reengineered, but that improved binding affinity alone is not sufficient to improve regulation. Instead, successful redesign requires preservation of the conformational dynamics that underlie physiological function. In our study, a variant optimized using static structural criteria increased affinity for both the RyR2 peptide and the intact channel. Yet this gain came at a cost: it distorted peptide geometry and worsened pathological Ca^2+^ leak. By contrast, a variant designed with dynamic structural constraints increased affinity while preserving conformational integrity, leading to improved RyR2 regulation *ex vivo*. Together, these findings show that preservation of conformational dynamics, rather than binding optimization alone, is the key requirement for successful CaM redesign.

RyR2 gating is governed by the integration of luminal^55^ and cytosolic^56^ cues that coordinate channel opening and closure (**Fig. 1A**). CaM functions as a cytosolic Ca^2+^-dependent modulator that remains associated with cardiac RyR from beat to beat, tuning channel activity in response to local Ca^2+^ signals^39,57^. Rather than dissociating during normal excitation-contraction coupling, CaM engages RyR2 through distinct domain-specific roles, with the N-domain contributing to regulation during channel inactivation and the C-domain serving as a stable anchoring element^18,58-61^. These dynamic features of CaM–RyR2 regulation make this system a particularly informative model for functionally guided redesign, because they link mechanistically defined conformational behavior to measurable structural, biochemical, and physiological outcomes. To examine how these regulatory roles constrain CaM redesign, we first applied a *static structure-*guided redesign approach to engineer RCaM1, introducing two N-domain substitutions (S38E and Q41E) predicted to enhance binding to the RyR2 CaM-binding peptide (**Fig. 2**). Consistent with this design rationale, RCaM1 exhibited increased affinity for both the isolated peptide and intact channel, driven primarily by slowed peptide dissociation through N-domain stabilization (**Fig. 3**). However, despite these favorable biochemical properties, RCaM1 paradoxically worsened RyR2 regulation *ex vivo*, increasing Ca^2+^ leak in permeabilized cardiomyocytes from both leaky (RyR2-S2814D) and wild-type mouse models (**Fig. 4, Supplementary Fig. S8**). This dissociation between affinity and function indicates that strengthened static interactions alone are insufficient to ensure effective physiological regulation.

Molecular dynamics simulations provided a mechanistic explanation for this paradox. Wild-type CaM preferentially sampled dynamically “annealed” conformations that maintained a straight, mechanically stable RyR2 peptide, a geometry likely required to constrain channel reopening. Such domain-level compaction and peptide stabilization are not apparent in crystallographic or cryo-EM snapshots but emerge in ensemble-based descriptions captured by molecular-dynamics simulations or NMR analyses^23,30,41,48,62,63^. Because RCaM1 was optimized against a static structural model that did not encode these dynamic constraints^64^, it frequently adopted an “unlocked” conformational state that induced bending of the RyR2 peptide at residue S3592 (**Figs. 2, 4**). Although RCaM1 strengthened binding, this peptide deformation disrupted conformational integrity and destabilized channel regulation, consistent with prior work in other CaM-target systems showing that CaM binding can be coupled to disorder-to-order transitions in the CaM-binding domain^65-68^.

By contrast, RCaM2 was designed using dynamically “annealed” wild-type CaM conformations as templates. This dynamic-structure redesign preserved peptide geometry while enhancing binding affinity, resulting in improved RyR2 regulation *ex vivo*, as evidenced by reduced pathological Ca^2+^ leak (**Fig. 4**). The functional relevance of peptide integrity at this site is supported by structural data: the region surrounding S3592 is unresolved in Ca^2+^-free RyR2-CaM cryo-EM structures^39^ (PDB ID: 6JI8), consistent with intrinsic flexibility, and exhibits pronounced bending in leaky RyR2 variants, including the PKA-phosphorylated RyR2-R420W mutant, even in the presence of Ca^2+^-bound CaM^56^ (PDB IDs:8UXL, 8UXM). In contrast, wild-type RyR2 structures in the presence of Ca^2+^ retain a straight peptide conformation^39^ (PDB IDs: 6JIY, 6JV2) (**Fig. 4C**). Together, these correlations link peptide deformation at S3592 to pathological Ca^2+^ leak and identify conformational instability within the CaM-binding region as a key determinant of dysregulated RyR2 activity. More broadly, these findings show that iterative redesign integrating static structural models with conformational dynamics can reveal mechanistic constraints on regulation that are not apparent from static structures alone^23,32,69,70^. In the context of pathological RyR2 leak, these findings also highlight the therapeutic potential of redesigning CaM to achieve functionally selective correction of channel regulation.

Beyond RyR2, this study establishes a broader principle for re-engineering conserved, multifunctional and dynamic proteins. Many regulatory proteins, like CaM, operate not through rigid lock-and-key interactions but by sampling conformational ensembles that shift across atomic, biochemical, and physiological scales^30,31,64^. Our findings therefore highlight that redesign strategies must preserve these ensemble dynamics while strengthening binding. This principle may extend to a wide range of CaM–target systems and to other regulatory proteins in which flexibility is intrinsic to function^23,32-34,70^. More broadly, the *dynamic-structure redesign* framework presented here provides a conceptual guide for dissecting and engineering complex protein–protein interaction networks. By integrating static structural information with molecular-dynamics simulations and functional validation, this approach captures how conformational integrity, domain stabilization, and affinity together determine regulatory outcomes. As such, it offers not only a path forward for rational protein engineering but also a general strategy for probing the dynamic principles that underlie protein–protein regulation across biological systems.

## MATERIALS AND METHODS

Full details of all experimental protocols are described in Supporting Information (SI) Appendix.

### In silico

Calmodulin–RyR2 complexes were modeled using Ca^2+^-CaM bound to RyR1 (PDB 2BCX)^71^ as a template and refined by I-TASSER homology modeling^72^ and Rosetta FastRelax^73^, with the later-released CaM–RyR2 structure (PDB 6Y4O)^43^ incorporated into the ensemble. Computational redesign was performed in OSPREY 3.0^44^ using the bbk* algorithm^74^, evaluating single-site substitutions across interface residues, followed by molecular-dynamics simulations in GROMACS^75^ with the AMBER99SB-ILDN force field^76^, standard equilibration protocols, LINCS constraints^77^, and binding energetics were evaluated using MM/PBSA calculations as described^78^, as implemented in g_mmpbsa^79,80^.

### In vitro

Recombinant CaM proteins were prepared as described^81,82^ and characterized by steady-state and stopped-flow fluorescence assays to determine Ca^2+^ sensitivity, kinetics, and RyR2 peptide affinity using established methods^82-84^. Binding to intact RyR2 in SR vesicles was quantified by a fluorescence-lifetime FRET competition assay^85-88^.

### Ex vivo

All animal procedures were approved by Institutional Animal Care and Use Committee of Vanderbilt University (animal protocol #M1600259-00), the Ohio State University (animal protocol 2010A00000117-R5), and Mississippi State University (animal protocol #25-466) in accordance with NIH guidelines. Ventricular myocytes were isolated^18^ from WT and RyR2-S2814D^89^ mice, and Ca^2+^ sparks/waves were recorded in permeabilized cells loaded with Fluo-4 under controlled Ca^2+^ buffering; spark parameters, mass, and leak were quantified^90^ using ImageJ SparkMaster and nonparametric statistical analysis.

## ACKNOWLEDGMENTS

This work was supported by National Institutes of Health (NIH) grants (5R01HL138579 awarded to S. Györke and J. P. Davis, R35GM142868 awarded to C. N. Johnson), and by an American Heart Association Predoctoral Fellowship (#25PRE1377151, awarded to V. Bogdanov).

## REFERENCES

(1) Carafoli E, Krebs J. Why Calcium? How Calcium Became the Best Communicator. J Biol Chem. 2016 Sep 30;291(40):20849–20857. doi: 10.1074/jbc.R116.735894. Epub 2016 Jul 26. PMID: 27462077; PMCID: PMC5076498.

(2) Carafoli E. Calcium signaling: a tale for all seasons. Proc Natl Acad Sci U S A. 2002 Feb 5;99(3):1115–22. doi: 10.1073/pnas.032427999. PMID: 11830654; PMCID: PMC122154.

(3) Princen K, Van Dooren T, van Gorsel M, Louros N, Yang X, Dumbacher M, Bastiaens I, Coupet K, Dupont S, Cuveliers E, Lauwers A, Laghmouchi M, Vanwelden T, Carmans S, Van Damme N, Duhamel H, Vansteenkiste S, Prerad J, Pipeleers K, Rodiers O, De Ridder L, Claes S, Busschots Y, Pringels L, Verhelst V, Debroux E, Brouwer M, Lievens S, Tavernier J, Farinelli M, Hughes-Asceri S, Voets M, Winderickx J, Wera S, de Wit J, Schymkowitz J, Rousseau F, Zetterberg H, Cummings JL, Annaert W, Cornelissen T, De Winter H, De Witte K, Fivaz M, Griffioen G. Pharmacological modulation of septins restores calcium homeostasis and is neuroprotective in models of Alzheimer’s disease. Science. 2024 May 31;384(6699):eadd6260. doi: 10.1126/science.add6260. Epub 2024 May 31. PMID: 38815015; PMCID: PMC11827694.

(4) Cherubini M, Lopez-Molina L, Gines S. Mitochondrial fission in Huntington’s disease mouse striatum disrupts ER-mitochondria contacts leading to disturbances in Ca2+ efflux and Reactive Oxygen Species (ROS) homeostasis. Neurobiol Dis. 2020 Mar;136:104741. doi: 10.1016/j.nbd.2020.104741. Epub 2020 Jan 10. PMID: 31931142.

(5) Tang TS, Slow E, Lupu V, Stavrovskaya IG, Sugimori M, Llinás R, Kristal BS, Hayden MR, Bezprozvanny I. Disturbed Ca2+ signaling and apoptosis of medium spiny neurons in Huntington’s disease. Proc Natl Acad Sci U S A. 2005 Feb 15;102(7):2602–7. doi: 10.1073/pnas.0409402102. Epub 2005 Feb 3. PMID: 15695335; PMCID: PMC548984.

(6) Zaichick SV, McGrath KM, Caraveo G. The role of Ca2+ signaling in Parkinson’s disease. Dis Model Mech. 2017 May 1;10(5):519–535. doi: 10.1242/dmm.028738. PMID: 28468938; PMCID: PMC5451174.

(7) Nademanee K, Singh BN. Control of cardiac arrhythmias by calcium antagonism. Ann N Y Acad Sci. 1988;522:536–52. doi: 10.1111/j.1749-6632.1988.tb33397.x. PMID: 3288060.

(8) Godfraind T. Discovery and Development of Calcium Channel Blockers. Front Pharmacol. 2017 May 29;8:286. doi: 10.3389/fphar.2017.00286. PMID: 28611661; PMCID: PMC5447095.

(9) Bers DM. Cardiac excitation-contraction coupling. Nature. 2002 Jan 10;415(6868):198–205. doi: 10.1038/415198a. PMID: 11805843.

(10) Zheng S, Wang X, Zhao D, Liu H, Hu Y. Calcium homeostasis and cancer: insights from endoplasmic reticulum-centered organelle communications. Trends Cell Biol. 2023 Apr;33(4):312–323. doi: 11.1016/j.tcb.2022.07.004. Epub 2022 Jul 29. PMID: 35915027.

(11) Peersen OB, Madsen TS, Falke JJ. Intermolecular tuning of calmodulin by target peptides and proteins: differential effects on Ca2+ binding and implications for kinase activation. Protein Sci. 1997 Apr;6(4):794–807. doi: 10.1002/pro.5560060406. PMID: 9098889; PMCID: PMC2144748.

(12) Wilson MA, Brunger AT. The 1.0 A crystal structure of Ca(2+)-bound calmodulin: an analysis of disorder and implications for functionally relevant plasticity. J Mol Biol. 2000 Sep 1;301(5):1237–56. doi: 10.1006/jmbi.2000.4029. PMID: 10966818.

(13) Bogdanov V, Mariangelo JIE, Soltisz AM, Sakuta G, Pokrass A, Beard C, Orengo BH, Kalinin R, Ulker A, Yunker B, Tikunova S, Thuma J, Xu X, Hund TJ, Veeraraghavan R, Davis JP, Györke S. Distinct intracellular spatiotemporal expression of Calmodulin genes underlies functional diversity of Calmodulin-dependent signalling in cardiac myocytes. Cardiovasc Res. 2025 Jul 8;121(7):1052–1065. doi: 10.1093/cvr/cvaf059. PMID: 40273382.

(14) Yap KL, Kim J, Truong K, Sherman M, Yuan T, Ikura M. Calmodulin target database. J Struct Funct Genomics. 2000;1(1):8–14. doi: 10.1023/a:1011320027914. PMID: 12836676.

(15) Berchtold MW, Villalobo A. The many faces of calmodulin in cell proliferation, programmed cell death, autophagy, and cancer. Biochim Biophys Acta. 2014 Feb;1843(2):398–435. doi: 10.1016/j.bbamcr.2013.10.021. Epub 2013 Nov 2. PMID: 24188867.

(16) O’Day DH, Huber RJ. Calmodulin binding proteins and neuroinflammation in multiple neurodegenerative diseases. BMC Neurosci. 2022 Mar 4;23(1):10. doi: 10.1186/s12868-022-00695-y. PMID: 35246032; PMCID: PMC8896083.

(17) Nassal D, Gratz D, Hund TJ. Challenges and Opportunities for Therapeutic Targeting of Calmodulin Kinase II in Heart. Front Pharmacol. 2020 Feb 5;11:35. doi: 10.3389/fphar.2020.00035. PMID: 32116711; PMCID: PMC7012788.

(18) Liu B, Walton SD, Ho HT, Belevych AE, Tikunova SB, Bonilla I, Shettigar V, Knollmann BC, Priori SG, Volpe P, Radwański PB, Davis JP, Györke S. Gene Transfer of Engineered Calmodulin Alleviates Ventricular Arrhythmias in a Calsequestrin-Associated Mouse Model of Catecholaminergic Polymorphic Ventricular Tachycardia. J Am Heart Assoc. 2018 May 2;7(10):e008155. doi: 10.1161/JAHA.117.008155. PMID: 29720499; PMCID: PMC6015318.

(19) Porto EM, Komor AC, Slaymaker IM, Yeo GW. Base editing: advances and therapeutic opportunities. Nat Rev Drug Discov. 2020 Dec;19(12):839–859. doi: 10.1038/s41573-020-0084-6. Epub 2020 Oct 19. PMID: 33077937; PMCID: PMC7721651.

(20) Geng G, Xu Y, Hu Z, Wang H, Chen X, Yuan W, Shu Y. Viral and non-viral vectors in gene therapy: current state and clinical perspectives. EBioMedicine. 2025 Aug;118:105834. doi: 10.1016/j.ebiom.2025.105834. Epub 2025 Jul 1. PMID: 40602323; PMCID: PMC12271757.

(21) Koh HY, Zheng Y, Yang M, Arora R, Webb GI, Pan S, Li L, Church G. AI-driven protein design. Nat Rev Bioeng. 2025 Sep 8; doi: 10.1038/s44222-025-00349-8

(22) Surpeta B, Sequeiros-Borja CE, Brezovsky J. Dynamics, a Powerful Component of Current and Future in Silico Approaches for Protein Design and Engineering. Int J Mol Sci. 2020 Apr 14;21(8):2713. doi: 10.3390/ijms21082713. PMID: 32295283; PMCID: PMC7215530.

(23) Guo AB, Akpinaroglu D, Kelly MJS, Kortemme T. Deep learning–guided design of dynamic proteins. Science. 2025 May 22;388(6749), eadr7094. doi: 10.1126/science.adr7094. PMID: 39071443; PMCID: PMC11275770

(24) Shifman JM, Mayo SL. Exploring the origins of binding specificity through the computational redesign of calmodulin. Proc Natl Acad Sci U S A. 2003 Nov 11;100(23):13274–9. doi: 10.1073/pnas.2234277100. Epub 2003 Nov 3. PMID: 14597710; PMCID: PMC263780.

(25) Shifman JM, Mayo SL. Modulating calmodulin binding specificity through computational protein design. J Mol Biol. 2002 Oct 25;323(3):417–23. doi: 10.1016/s0022-2836(02)00881-1. PMID: 12381298.

(26) Yosef E, Politi R, Choi MH, Shifman JM. Computational design of calmodulin mutants with up to 900-fold increase in binding specificity. J Mol Biol. 2009 Feb 6;385(5):1470–80. doi: 10.1016/j.jmb.2008.09.053. Epub 2008 Sep 27. PMID: 18845160.

(27) Palmer AE, Giacomello M, Kortemme T, Hires SA, Lev-Ram V, Baker D, Tsien RY. Ca2+ indicators based on computationally redesigned calmodulin-peptide pairs. Chem Biol. 2006 May;13(5):521–30. doi: 10.1016/j.chembiol.2006.03.007. PMID: 16720273.

(28) Fischer R, Koller M, Flura M, Mathews S, Strehler-Page MA, Krebs J, Penniston JT, Carafoli E, Strehler EE. Multiple divergent mRNAs code for a single human calmodulin. J Biol Chem. 1988 Nov 15;263(32):17055–62. PMID: 3182832.

(29) Yang T, Poovaiah BW. Calcium/calmodulin-mediated signal network in plants. Trends Plant Sci. 2003 Oct;8(10):505–12. doi: 10.1016/j.tplants.2003.09.004. PMID: 14557048.

(30) Boehr DD, Nussinov R, Wright PE. The role of dynamic conformational ensembles in biomolecular recognition. Nat Chem Biol. 2009 Nov;5(11):789–96. doi: 10.1038/nchembio.232. Erratum in: Nat Chem Biol. 2009 Dec;5(12):954. PMID: 19841628; PMCID: PMC2916928.

(31) Henzler-Wildman K, Kern D. Dynamic personalities of proteins. Nature. 2007 Dec 13;450(7172):964–72. doi: 10.1038/nature06522. PMID: 18075575.

(32) Sauer MF, Sevy AM, Crowe JE Jr, Meiler J. Multi-state design of flexible proteins predicts sequences optimal for conformational change. PLoS Comput Biol. 2020 Feb 7;16(2):e1007339. doi: 10.1371/journal.pcbi.1007339. PMID: 32032348; PMCID: PMC7032724.

(33) Davey JA, Chica RA. Multistate approaches in computational protein design. Protein Sci. 2012 Sep;21(9):1241–52. doi: 10.1002/pro.2128. Epub 2012 Aug 10. PMID: 22811394; PMCID: PMC3631354.

(34) Lassila JK. Conformational diversity and computational enzyme design. Curr Opin Chem Biol. 2010 Oct;14(5):676–82. doi: 10.1016/j.cbpa.2010.08.010. Epub 2010 Sep 7. PMID: 20829099; PMCID: PMC2953567.

(35) Zheng J, Ponce-Balbuena D, Ríos Pérez EB, Xiao L, Dooge HC, Valdivia HH, Alvarado FJ. Preventing the phosphorylation of RyR2 at canonical sites reduces Ca2+ leak and promotes arrhythmia by reactivating the INa current. Nat Cardiovasc Res. 2025 Aug;4(8):976–990. doi: 10.1038/s44161-025-00693-3. Epub 2025 Aug 12. PMID: 40797044; PMCID: PMC12343298.

(36) Yamaguchi N, Xu L, Pasek DA, Evans KE, Meissner G. Molecular basis of calmodulin binding to cardiac muscle Ca(2+) release channel (ryanodine receptor). J Biol Chem. 2003 Jun 27;278(26):23480–6. doi: 10.1074/jbc.M301125200. Epub 2003 Apr 21. PMID: 12707260.

(37) Xu X, Yano M, Uchinoumi H, Hino A, Suetomi T, Ono M, Tateishi H, Oda T, Okuda S, Doi M, Kobayashi S, Yamamoto T, Ikeda Y, Ikemoto N, Matsuzaki M. Defective calmodulin binding to the cardiac ryanodine receptor plays a key role in CPVT-associated channel dysfunction. Biochem Biophys Res Commun. 2010 Apr 9;394(3):660–6. doi: 10.1016/j.bbrc.2010.03.046. Epub 2010 Mar 10. PMID: 20226167; PMCID: PMC2858291.

(38) Belevych AE, Radwański PB, Carnes CA, Györke S. ‘Ryanopathy’: causes and manifestations of RyR2 dysfunction in heart failure. Cardiovasc Res. 2013 May 1;98(2):240–7. doi: 10.1093/cvr/cvt024. Epub 2013 Feb 12. PMID: 23408344; PMCID: PMC3633158.

(39) Gong D, Chi X, Wei J, Zhou G, Huang G, Zhang L, Wang R, Lei J, Chen SRW, Yan N. Modulation of cardiac ryanodine receptor 2 by calmodulin. Nature. 2019 Aug;572(7769):347–351. doi: 10.1038/s41586-019-1377-y. Epub 2019 Jul 5. PMID: 31278385.

(40) Rebbeck RT, Svensson B, Zhang J, Samsó M, Thomas DD, Bers DM, Cornea RL. Kinetics and mapping of Ca-driven calmodulin conformations on skeletal and cardiac muscle ryanodine receptors. Nat Commun. 2024 Jun 15;15(1):5120. doi: 10.1038/s41467-024-48951-5. PMID: 38879623; PMCID: PMC11180167.

(41) Cui X, Ge L, Chen X, Lv Z, Wang S, Zhou X, Zhang G. Beyond static structures: protein dynamic conformations modeling in the post-AlphaFold era. Brief Bioinform. 2025 Jul 2;26(4):bbaf340. doi: 10.1093/bib/bbaf340. PMID: 40663654; PMCID: PMC12262120.

(42) Davis JP, Shettigar V, Tikunova SB, Little SC, Liu B, Siddiqui JK, Janssen PM, Ziolo MT, Walton SD. Designing proteins to combat disease: Cardiac troponin C as an example. Arch Biochem Biophys. 2016 Jul 1;601:4–10. doi: 10.1016/j.abb.2016.02.007. Epub 2016 Feb 18. PMID: 26901433; PMCID: PMC4899299.

(43) Holt C, Hamborg L, Lau K, Brohus M, Sørensen AB, Larsen KT, Sommer C, Van Petegem F, Overgaard MT, Wimmer R. The arrhythmogenic N53I variant subtly changes the structure and dynamics in the calmodulin N-terminal domain, altering its interaction with the cardiac ryanodine receptor. J Biol Chem. 2020 May 29;295(22):7620–7634. doi: 10.1074/jbc.RA120.013430. Epub 2020 Apr 21. PMID: 32317284; PMCID: PMC7261784.

(44) Hallen MA, Martin JW, Ojewole A, Jou JD, Lowegard AU, Frenkel MS, Gainza P, Nisonoff HM, Mukund A, Wang S, Holt GT, Zhou D, Dowd E, Donald BR. OSPREY 3.0: Open-source protein redesign for you, with powerful new features. J Comput Chem. 2018 Nov 15;39(30):2494–2507. doi: 10.1002/jcc.25522. Epub 2018 Oct 14. PMID: 30368845; PMCID: PMC6391056.

(45) Mandell DJ, Kortemme T. Backbone flexibility in computational protein design. Curr Opin Biotechnol. 2009 Aug;20(4):420–8. doi: 10.1016/j.copbio.2009.07.006. Epub 2009 Aug 24. PMID: 19709874.

(46) Davis JP, Tikunova SB, Walsh MP, Johnson JD. Characterizing the response of calcium signal transducers to generated calcium transients. Biochemistry. 1999 Mar 30;38(13):4235–44. doi: 10.1021/bi982495z. PMID: 10194340.

(47) Kumari R, Kumar R; Open Source Drug Discovery Consortium; Lynn A. g_mmpbsa--a GROMACS tool for high-throughput MM-PBSA calculations. J Chem Inf Model. 2014 Jul 28;54(7):1951–62. doi: 10.1021/ci500020m. Epub 2014 Jun 19. PMID: 24850022.

(48) Childers MC, Daggett V. Insights from molecular dynamics simulations for computational protein design. Mol Syst Des Eng. 2017 Feb 1;2(1):9–33. doi: 10.1039/C6ME00083E. Epub 2017 Jan 9. PMID: 28239489; PMCID: PMC5321087.

(49) Jarmoskaite I, AlSadhan I, Vaidyanathan PP, Herschlag D. How to measure and evaluate binding affinities. Elife. 2020 Aug 6;9:e57264. doi: 10.7554/eLife.57264. PMID: 32758356; PMCID: PMC7452723.

(50) Davis JP, Tikunova SB, Walsh MP, Johnson JD. Characterizing the response of calcium signal transducers to generated calcium transients. Biochemistry. 1999 Mar 30;38(13):4235–44. doi: 10.1021/bi982495z. PMID: 10194340.

(51) Van Lierop JE, Wilson DP, Davis JP, Tikunova S, Sutherland C, Walsh MP, Johnson JD. Activation of smooth muscle myosin light chain kinase by calmodulin. Role of LYS(30) and GLY(40). J Biol Chem. 2002 Feb 22;277(8):6550–8. doi: 10.1074/jbc.M111404200. Epub 2001 Dec 17. PMID: 11748245.

(52) Cheng H, Lederer MR, Lederer WJ, Cannell MB. Calcium sparks and [Ca2+]i waves in cardiac myocytes. Am J Physiol. 1996 Jan;270(1 Pt 1):C148–59. doi: 10.1152/ajpcell.1996.270.1.C148. PMID: 8772440.

(53) Radwański PB, Belevych AE, Brunello L, Carnes CA, Györke S. Store-dependent deactivation: cooling the chain-reaction of myocardial calcium signaling. J Mol Cell Cardiol. 2013 May;58:77–83. doi: 10.1016/j.yjmcc.2012.10.008. Epub 2012 Oct 27. PMID: 23108187; PMCID: PMC4068615.

(54) Marx SO, Reiken S, Hisamatsu Y, Jayaraman T, Burkhoff D, Rosemblit N, Marks AR. PKA phosphorylation dissociates FKBP12.6 from the calcium release channel (ryanodine receptor): defective regulation in failing hearts. Cell. 2000 May 12;101(4):365–76. doi: 10.1016/s0092-8674(00)80847-8. PMID: 10830164.

(55) Terentyev D, Viatchenko-Karpinski S, Valdivia HH, Escobar AL, Györke S. Luminal Ca2+ controls termination and refractory behavior of Ca2+-induced Ca2+ release in cardiac myocytes. Circ Res. 2002 Sep 6;91(5):414–20. doi: 10.1161/01.res.0000032490.04207.bd. PMID: 12215490.

(56) Miotto MC, Reiken S, Wronska A, Yuan Q, Dridi H, Liu Y, Weninger G, Tchagou C, Marks AR. Structural basis for ryanodine receptor type 2 leak in heart failure and arrhythmogenic disorders. Nat Commun. 2024 Sep 15;15(1):8080. doi: 10.1038/s41467-024-51791-y. PMID: 39278969; PMCID: PMC11402997.

(57) Xu L, Meissner G. Mechanism of calmodulin inhibition of cardiac sarcoplasmic reticulum Ca2+ release channel (ryanodine receptor). Biophys J. 2004 Feb;86(2):797–804. doi: 10.1016/S0006-3495(04)74155-7. PMID: 14747315; PMCID: PMC1303927.

(58) Wu X, Bers DM. Free and bound intracellular calmodulin measurements in cardiac myocytes. Cell Calcium. 2007 Apr;41(4):353–64. doi: 10.1016/j.ceca.2006.07.011. Epub 2006 Sep 26. PMID: 16999996; PMCID: PMC1868497.

(59) Ono M, Yano M, Hino A, Suetomi T, Xu X, Susa T, Uchinoumi H, Tateishi H, Oda T, Okuda S, Doi M, Kobayashi S, Yamamoto T, Koseki N, Kyushiki H, Ikemoto N, Matsuzaki M. Dissociation of calmodulin from cardiac ryanodine receptor causes aberrant Ca(2+) release in heart failure. Cardiovasc Res. 2010 Sep 1;87(4):609–17. doi: 10.1093/cvr/cvq108. Epub 2010 Apr 13. PMID: 20388639; PMCID: PMC3025723.

(60) Guo T, Fruen BR, Nitu FR, Nguyen TD, Yang Y, Cornea RL, Bers DM. FRET detection of calmodulin binding to the cardiac RyR2 calcium release channel. Biophys J. 2011 Nov 2;101(9):2170–7. doi: 10.1016/j.bpj.2011.09.030. Epub 2011 Nov 1. PMID: 22067155; PMCID: PMC3207159.

(61) Yang Y, Guo T, Oda T, Chakraborty A, Chen L, Uchinoumi H, Knowlton AA, Fruen BR, Cornea RL, Meissner G, Bers DM. Cardiac myocyte Z-line calmodulin is mainly RyR2-bound, and reduction is arrhythmogenic and occurs in heart failure. Circ Res. 2014 Jan 17;114(2):295–306. doi: 10.1161/CIRCRESAHA.114.302857. Epub 2013 Nov 1. PMID: 24186966; PMCID: PMC4004530.

(62) Robustelli P, Stafford KA, Palmer AG 3rd. Interpreting protein structural dynamics from NMR chemical shifts. J Am Chem Soc. 2012 Apr 11;134(14):6365–74. doi: 10.1021/ja300265w. Epub 2012 Mar 28. PMID: 22381384; PMCID: PMC3324661.

(63) Lewis S, Hempel T, Jiménez-Luna J, Gastegger M, Xie Y, Foong AYK, Satorras VG, Abdin O, Veeling BS, Zaporozhets I, Chen Y, Yang S, Foster AE, Schneuing A, Nigam J, Barbero F, Stimper V, Campbell A, Yim J, Lienen M, Shi Y, Zheng S, Schulz H, Munir U, Sordillo R, Tomioka R, Clementi C, Noé F. Scalable emulation of protein equilibrium ensembles with generative deep learning. Science. 2025 Aug 14;389(6761):eadv9817. doi: 10.1126/science.adv9817. Epub 2025 Aug 14. PMID: 40638710.

(64) Rennella E, Sahtoe DD, Baker D, Kay LE. Exploiting conformational dynamics to modulate the function of designed proteins. Proc Natl Acad Sci U S A. 2023 May 2;120(18):e2303149120. doi: 10.1073/pnas.2303149120. Epub 2023 Apr 24. PMID: 37094170; PMCID: PMC10161014.

(65) O’Neil KT, DeGrado WF. How calmodulin binds its targets: sequence independent recognition of amphiphilic alpha-helices. Trends Biochem Sci. 1990 Feb;15(2):59–64. doi: 10.1016/0968-0004(90)90177-d. PMID: 2186516.

(66) Ikura M, Clore GM, Gronenborn AM, Zhu G, Klee CB, Bax A. Solution structure of a calmodulin-target peptide complex by multidimensional NMR. Science. 1992 May 1;256(5057):632–8. doi: 10.1126/science.1585175. PMID: 1585175.

(67) Zhang Y, Tan H, Chen G, Jia Z. Investigating the disorder-order transition of calmodulin binding domain upon binding calmodulin using molecular dynamics simulation. J Mol Recognit. 2010 Jul-Aug;23(4):360–8. doi: 10.1002/jmr.1002. PMID: 19998355.

(68) Zhang M, Vogel HJ. The calmodulin-binding domain of caldesmon binds to calmodulin in an alpha-helical conformation. Biochemistry. 1994 Feb 8;33(5):1163–71. doi: 10.1021/bi00171a016. PMID: 8110748.

(69) Praetorius F, Leung PJY, Tessmer MH, Broerman A, Demakis C, Dishman AF, Pillai A, Idris A, Juergens D, Dauparas J, Li X, Levine PM, Lamb M, Ballard RK, Gerben SR, Nguyen H, Kang A, Sankaran B, Bera AK, Volkman BF, Nivala J, Stoll S, Baker D. Design of stimulus-responsive two-state hinge proteins. Science. 2023 Aug 18;381(6659):754–760. doi: 10.1126/science.adg7731. Epub 2023 Aug 17. PMID: 37590357; PMCID: PMC10697137.

(70) Jefferson RE, Oggier A, Füglistaler A, Camviel N, Hijazi M, Villarreal AR, Arber C, Barth P. Computational design of dynamic receptor-peptide signaling complexes applied to chemotaxis. Nat Commun. 2023 May 19;14(1):2875. doi: 10.1038/s41467-023-38491-9. PMID: 37208363; PMCID: PMC10198977.

(71) Maximciuc AA, Putkey JA, Shamoo Y, Mackenzie KR. Complex of calmodulin with a ryanodine receptor target reveals a novel, flexible binding mode. Structure. 2006 Oct;14(10):1547–56. doi: 10.1016/j.str.2006.08.011. PMID: 17027503.

(72) Yang J, Zhang Y. I-TASSER server: new development for protein structure and function predictions. Nucleic Acids Res. 2015 Jul 1;43(W1):W174–81. doi: 10.1093/nar/gkv342. Epub 2015 Apr 16. PMID: 25883148; PMCID: PMC4489253.

(73) Adolf-Bryfogle J, Kalyuzhniy O, Kubitz M, Weitzner BD, Hu X, Adachi Y, Schief WR, Dunbrack RL Jr. RosettaAntibodyDesign (RAbD): A general framework for computational antibody design. PLoS Comput Biol. 2018 Apr 27;14(4):e1006112. doi: 10.1371/journal.pcbi.1006112. PMID: 29702641; PMCID: PMC5942852.

(74) Ojewole AA, Jou JD, Fowler VG, Donald BR. BBK* (Branch and Bound Over K*): A Provable and Efficient Ensemble-Based Protein Design Algorithm to Optimize Stability and Binding Affinity Over Large Sequence Spaces. J Comput Biol. 2018 Jul;25(7):726–739. doi: 10.1089/cmb.2017.0267. Epub 2018 Mar 13. PMID: 29641249; PMCID: PMC6074059.

(75) Abraham M, Alekseenko A; Bergh C; Blau C; Briand E; Doijade M; Fleischmann S; Gapsys V; Garg G; Gorelov S; Gouaillardet G; Gray A; Irrgang ME; Jalalypour F; Jordan J; Junghans C; Kanduri P; Keller S; Kutzner C; Lemkul JA; Lundborg M; Merz P; Miletic V; Morozov D; Páll S; Schulz R; Shirts M; Shvetsov A; Soproni B; van der Spoel D; Turner P; Uphoff C; Villa A; Wingbermühle S; Zhmurov A; Bauer P; Hess B; Lindahl E. GROMACS 2023.1 Manual (2023.1). Zenodo. 10.5281/zenodo.7852189

(76) Lindorff-Larsen K, Piana S, Palmo K, Maragakis P, Klepeis JL, Dror RO, Shaw DE. Improved side-chain torsion potentials for the Amber ff99SB protein force field. Proteins. 2010 Jun;78(8):1950–8. doi: 10.1002/prot.22711. PMID: 20408171; PMCID: PMC2970904.

(77) Hess B, Bekker H, Berendsen HJ. Fraaije JGEM. LINCS: A linear constraint solver for molecular simulations. J Comput Chem. 1997;18(12):1463–72.

(78) Scott CE, Kekenes-Huskey PM. Molecular Basis of S100A1 Activation at Saturating and Subsaturating Calcium Concentrations. Biophys J. 2016 Mar 8;110(5):1052–63. doi: 10.1016/j.bpj.2015.12.040. Erratum in: Biophys J. 2016 Apr 26;110(8):1907. doi: 10.1016/j.bpj.2016.03.032. PMID: 26958883; PMCID: PMC4788715.

(79) Kumari R, Kumar R; Open Source Drug Discovery Consortium; Lynn A. g_mmpbsa--a GROMACS tool for high-throughput MM-PBSA calculations. J Chem Inf Model. 2014 Jul 28;54(7):1951–62. doi: 10.1021/ci500020m. Epub 2014 Jun 19. PMID: 24850022.

(80) Baker NA, Sept D, Joseph S, Holst MJ, McCammon JA. Electrostatics of nanosystems: application to microtubules and the ribosome. Proc Natl Acad Sci U S A. 2001 Aug 28;98(18):10037–41. doi: 10.1073/pnas.181342398. Epub 2001 Aug 21. PMID: 11517324; PMCID: PMC56910.

(81) Walton SD, Chakravarthy H, Shettigar V, O’Neil AJ, Siddiqui JK, Jones BR, Tikunova SB, Davis JP. Divergent Soybean Calmodulins Respond Similarly to Calcium Transients: Insight into Differential Target Regulation. Front Plant Sci. 2017 Feb 15;8:208. doi: 10.3389/fpls.2017.00208. PMID: 28261258; PMCID: PMC5309217.

(82) Black DJ, Tikunova SB, Johnson JD, Davis JP. Acid pairs increase the N-terminal Ca2+ affinity of CaM by increasing the rate of Ca2+ association. Biochemistry. 2000 Nov 14;39(45):13831–7. doi: 10.1021/bi001106+. PMID: 11076523.

(83) Robertson S, Potter JD. The regulation of free Ca2+ion concentration by metal chelators. In: Schwartz A, editor. Myocardial Biology. Boston, MA: Springer US; 1984. p. 63–75. doi: 10.1007/978-1-4684-4778-1_6.

(84) Tikunova SB, Rall JA, Davis JP. Effect of hydrophobic residue substitutions with glutamine on Ca(2+) binding and exchange with the N-domain of troponin C. Biochemistry. 2002 May 28;41(21):6697–705. doi: 10.1021/bi011763h. PMID: 12022873.

(85) Hwang HS, Nitu FR, Yang Y, Walweel K, Pereira L, Johnson CN, Faggioni M, Chazin WJ, Laver D, George AL Jr, Cornea RL, Bers DM, Knollmann BC. Divergent regulation of ryanodine receptor 2 calcium release channels by arrhythmogenic human calmodulin missense mutants. Circ Res. 2014 Mar 28;114(7):1114–24. doi: 10.1161/CIRCRESAHA.114.303391. Epub 2014 Feb 21. PMID: 24563457; PMCID: PMC3990285.

(86) Fruen BR, Bardy JM, Byrem TM, Strasburg GM, Louis CF. Differential Ca(2+) sensitivity of skeletal and cardiac muscle ryanodine receptors in the presence of calmodulin. Am J Physiol Cell Physiol. 2000 Sep;279(3):C724–33. doi: 10.1152/ajpcell.2000.279.3.C724. PMID: 10942723.

(87) Petersen KJ, Peterson KC, Muretta JM, Higgins SE, Gillispie GD, Thomas DD. Fluorescence lifetime plate reader: resolution and precision meet high-throughput. Rev Sci Instrum. 2014 Nov;85(11):113101. doi: 10.1063/1.4900727. PMID: 25430092; PMCID: PMC4242087.

(88) Rebbeck RT, Essawy MM, Nitu FR, Grant BD, Gillispie GD, Thomas DD, Bers DM, Cornea RL. High-Throughput Screens to Discover Small-Molecule Modulators of Ryanodine Receptor Calcium Release Channels. SLAS Discov. 2017 Feb;22(2):176–186. doi: 10.1177/1087057116674312. Epub 2016 Oct 22. PMID: 27760856; PMCID: PMC5337111.

(89) van Oort RJ, McCauley MD, Dixit SS, Pereira L, Yang Y, Respress JL, Wang Q, De Almeida AC, Skapura DG, Anderson ME, Bers DM, Wehrens XH. Ryanodine receptor phosphorylation by calcium/calmodulin-dependent protein kinase II promotes life-threatening ventricular arrhythmias in mice with heart failure. Circulation. 2010 Dec 21;122(25):2669–79. doi: 10.1161/CIRCULATIONAHA.110.982298. Epub 2010 Nov 15. PMID: 21098440; PMCID: PMC3075419.

(90) Hollingworth S, Peet J, Chandler WK, Baylor SM. Calcium sparks in intact skeletal muscle fibers of the frog. J Gen Physiol. 2001 Dec;118(6):653–78. doi: 10.1085/jgp.118.6.653. PMID: 11723160; PMCID: PMC2229509.

