## SUPPORTING INFORMATION for "Dynamic-Structure Redesign of Calmodulin Reveals Mechanistic Constraints on Ryr2 Regulation"

**SUPPORTING INFORMATION (SI) APPENDIX**

This PDF file includes:

**1. TITLE PAGE**
**2. MATERIALS AND METHODS**
**3. SUPPLEMENTARY FIGURES** (Figures S1-S8)
**4. SUPPLEMENTARY MOVIE LEGEND**

#### 1. TITLE PAGE

### Dynamic-Structure Redesign of Calmodulin Reveals Mechanistic Constraints on Ryr2 Regulation

Vladimir Bogdanov[1,2], Svetlana Tikunova[1,2], Nicolas Fadell[2], Robyn T. Rebbeck[3], Melanie L. Aprahamian[4], Md Nure Alam Afsar[5], Aleksei Chekodanov[6], Daniel J. Blackwell[7], Bjorn C. Knollmann[7], Razvan L. Cornea[3], Pete M. Kekenes-Huskey[8], Steffen Lindert[9], Christopher N. Johnson[5], Sandor Györke*[1,2] & Jonathan P. Davis*[1,2]

[1]The Dorothy M. Davis Heart and Lung Research Institute, The Ohio State University Wexner Medical Center, Columbus, Ohio, 43210
[2]Department of Physiology and Cell Biology, College of Medicine, The Ohio State University, Columbus, Ohio, 43210
[3]Department of Biochemistry, Molecular Biology and Biophysics, University of Minnesota, Minnesota, Minneapolis, 55455
[4]Department of Chemistry and Biochemistry, Ohio State University, Columbus, Ohio 43210, United States
[5]Department of Chemistry, Mississippi State University, Starkville, Mississippi, 39759
[6]Independent Software Developer, Moscow, Russia, 119049.
[7]Vanderbilt Center for Arrhythmia Research and Therapeutics, Division of Clinical Pharmacology, Vanderbilt University Medical Center, Nashville, Tennessee, 37212
[8]Department of Cell and Molecular Physiology, Stritch School of Medicine, Loyola University Chicago, Maywood, Illinois, 60153
[9]Department of Chemistry and Biochemistry, The University of California, Los Angeles, California, 90095

**Corresponding author:**

**Author Contributions:** Conceptualization, J.P.D. and V.B.; Methodology, J.P.D., V.B., R.L.C., P.M.K-H., S.L., C.N.J., and S.G.; Software, A.C., V.B.; Validation, J.P.D., V.B., R.L.C., P.M.K-H., S.L., C.N.J., and S.T.; Formal Analysis, J.P.D., V.B., S.T., R.T.R., M.L.A., M.N.A, C.N.J., and S.L.; Investigation, J.P.D., V.B., S.T., R.T.R., M.L.A., M.N.A, C.N.J., and S.L.; Resources, J.P.D., S.G., R.L.C., C.N.J., D.J.B., B.C.K., and S.L.; Data Curation, V.B. and J.P.D.; Writing - Original Draft, J.P.D., V.B., M.N.A, and C.N.J; Writing – Review and Editing, J.P.D., V.B., R.L.C., P.M.K-H., S.L., C.N.J., S.T., and S.G.; Visualization, V.B., J.P.D., R.T.R., and C.N.J.; Supervision, J.P.D., V.B., S.G. and C.N.J.; Funding Acquisition, J.P.D., S.G., R.L.C., S.L., C.N.J. and P.M.K-H.

**Competing Interest Statement:** The authors declare no competing interests.

**Disclaimer**
Razvan Cornea is currently an employee of the National Institutes of Health. This work was conducted during his previous employment, at University of Minnesota – Twin Cities. The opinions expressed in this article are the author's own and do not reflect the view of the National Institutes of Health, the Department of Health and Human Services, or the United States government.

**Classification:**
Major: Biological Sciences
Minor: Biophysics and Computational Biology

**Keywords:**
calmodulin; ryanodine receptor; calcium signaling; molecular dynamics simulations; protein engineering; cardiac physiology

#### 2. MATERIALS AND METHODS

##### *Materials*

Phenyl-Sepharose CL-4B, EGTA, Quin-2 and Liberase TH were purchased from Millipore Sigma (St. Loius, MO, USA). Fluo-4 was purchased from Thermo Fisher Scientific (Waltham, MA, USA). All other chemicals were purchased from Fisher Scientific (Pittsburg, PA, USA). The CaM-binding domain of human RYR2 receptor (RSKKAVWHKLLSKQRKRAVVACFRMAPLYNL) was synthesized by Ohio Peptide LLC (Powell, OH, USA).

##### *CaM-RyR2 peptide complex modeling*

We began with the $Ca^{2+}$-bound CaM–RyR1 peptide crystal structure (PDB ID: 2BCX)[71] as an initial template. The RyR1 peptide was converted in silico to the corresponding RyR2 sequence (K3579R, R3594K, T3604A) and renumbered to match the human RyR2 peptide spanning residues 3581–3612. The full-length 147-residue CaM was then modeled in its appropriate N- and C-domain conformations using I-TASSER homology modeling[72]. For each CaM–RyR2 complex, 2,000 models were generated using Rosetta FastRelax[73], and the top eight low-energy structures were selected as the relaxed structural ensemble used for subsequent OSPREY redesign calculations.

During the course of this work, a high-resolution structure of $Ca^{2+}$–CaM bound to the RyR2 3581–3612 peptide (PDB ID: 6Y4O)[43] became available. This structure was incorporated as an additional template (**Fig. 2A, model 9**) and processed identically to the FastRelax-generated models for redesign analysis.

##### *Computational Protein Redesign*

In the relaxed CaM–RyR2 complexes, CaM residues within 7 Å of the RyR2 peptide (distance measured after hydrogen placement and structural relaxation) were identified using a custom Python workflow (hyperlink: Residues Identification). All redesign calculations were carried out using the complete human CaM sequence (147 amino acids) bound to the RyR2 CaM-binding segment spanning residues 3581–3612. These residues, together with their immediately adjacent positions, were designated as design sites. All redesign calculations were performed using OSPREY 3.0[44], employing the bbk* algorithm[74]. At each design position, only a single CaM residue was permitted to mutate, whereas neighboring residues on both CaM (protein strand) and the RyR2 peptide (ligand strand) were modeled as flexible. Flexibility was restricted to wild-type rotamers with continuous side-chain sampling, allowing local structural relaxation without introducing additional mutations. No multi-site or combinatorial mutations were evaluated (hyperlink: OSPREY Preprocessing).

All bbk* computations were carried out using batch script (Run OSPREY Batch) on a Linux workstation equipped with a 32-core AMD 3970X Threadripper CPU, an NVIDIA RTX 3090 GPU, and 128 GB system RAM. OSPREY was run through Python 3.9 and Java 14 (OSPREY 3.0) with a Java heap size of 100,000 MiB (8,192 MiB reserved for garbage collection and ~91,808 MiB available for computation). Within OSPREY, parallelism was configured to use 32 CPU cores and one CUDA-enabled GPU with 82 streams. Energy evaluations used OSPREY's default AMBER-based molecular mechanics force field (ff99/ff14SB hybrid with Lazaridis–Karplus implicit solvation). Input PDB files were standardized to AMBER residue-naming conventions to ensure compatibility with the template library. Bbk* calculations used an ε-tolerance of 0.01.

Across all relaxed CaM–RyR2 structures, each design position was independently subjected to single-site redesign against all 20 canonical amino acids. For every wild-type and mutant sequence, bbk* computed a K* score. $\Delta k^*_{score}$ values were then calculated in post-processing (available via hyperlink: TSV Processing), according to:

$$\Delta k^*_{score} = k^*_{score\ mutant} - k^*_{score\ wt}$$

Higher $\Delta k^*_{score}$ values correspond to more preferred substitutions. The resulting rankings are presented in **Figure 1A, C**. CaM–RyR2 complexes incorporating top-ranked substitutions were subsequently used for molecular-dynamics simulations.

***Molecular dynamics Simulations***

RCaM-RyR2 complexes were parameterized using the AMBER99SB-ILDN force field[76] with GROMACS 2018.5 software[75]. The complexes were solvated in a TIP3P water model and placed into a simulation box, with a minimum distance of 10 Å from the protein to the box edge. The solvated complexes were neutralized with $K^+$ and $Cl^-$ ions, replacing water molecules to achieve a physiological condition of a total concentration of 0.15 M KCl. The neutralized systems were minimized until the maximum force was less than 1000.0 $kJ \cdot mol^{-1} \cdot nm^{-1}$, allowing up to 50000 steps. Long-range electrostatic interactions were treated using the Particle Mesh Ewald (PME) method. A cut-off of 1.2 nm was used for both short-range electrostatic and Van der Waals interactions. Periodic boundary conditions (PBC) were applied in all three dimensions. Following energy minimization, equilibration was carried out in two stages: a constant number, volume, and temperature (NVT) ensemble, and a constant number, pressure, and temperature (NPT) ensemble. Bonds involving hydrogen atoms were constrained using the LINear Constraint Solver (LINCS)[77] allowing a 2-fs integration timestep. The system was gradually heated to 297.15 K using initial velocities generated from a Maxwell-Boltzmann distribution at the same temperature and then maintained using the V-rescale thermostat. Pressure coupling was not applied during the NVT stage. During the NPT equilibration, the Parrinello-Rahman barostat was used to maintain a reference pressure of 1.0 bar with an isothermal compressibility of $4.5 \cdot 10^{-5}$ $bar^{-1}$. A molecular dynamics production run of at least 100 ns was carried out under the same NPT conditions as the second equilibration phase. The coordinates, velocities, and energies were saved every 10 ps for further analysis. No initial velocities were generated for this phase, as it was a continuation from the NPT equilibration.

***MM/PBSA analysis***

Binding free energies were estimated from molecular dynamics trajectories using the Molecular Mechanics/Poisson–Boltzmann Surface Area (MM/PBSA) method as implemented in g_mmpbsa[79,80]. Trajectories were re-centered to the primary periodic image prior to analysis, and snapshots were extracted every 1 ns from the final 100 ns of each production trajectory.

The binding free energy was calculated as:

$$\Delta G_{\text{bind}} = \langle G_{\text{complex}} \rangle - \langle G_{\text{receptor}} \rangle - \langle G_{\text{ligand}} \rangle$$

The free energy terms comprise molecular mechanics (van der Waals and electrostatic) and solvation contributions calculated using Poisson–Boltzmann electrostatics (solute dielectric = 8; solvent dielectric = 80) and a solvent-accessible surface area–based nonpolar model, following the default g_mmpbsa protocol. Entropic contributions were not included.

Because MM/PBSA calculations are most reliable for relative comparisons rather than absolute free energy estimation, the resulting $\Delta G_{\text{bind}}$ values were used to rank binding affinities and identify energetic trends rather than to infer absolute thermodynamic quantities. Reported MM/PBSA energy components correspond to the standard g_mmpbsa output terms ($\Delta E_{\text{MM}}$, $\Delta G_{\text{polar}}$, $\Delta G_{\text{nonpolar}}$, and $\Delta G_{\text{bind}}$) and were averaged over all analyzed snapshots. Per-residue free energy decomposition was performed using the built-in g_mmpbsa decomposition scheme to identify residues contributing most strongly to binding. Graphical visualization was performed using Grace.

***Preparation of Proteins.***

The recombinant wild-type vertebrate CaM (wtCaM) and its mutants were generated, expressed and purified via standard laboratory techniques, as previously described[81,82].

***Steady-State Fluorescence Measurements.***

All steady-state fluorescence measurements were carried out using a Perkin-Elmer LS55 fluorescence spectrometer at 20°C. Steady-state experiments were carried out in titration buffer (200 mM MOPS (to prevent pH changes upon addition of $Ca^{2+}$), 150 mM KCl, 2 mM EGTA, pH 7.0) at 20°C. In order to follow the $Ca^{2+}$-dependent conformational change in the C-domain of CaM in the absence of RyR2 peptide, Tyr fluorescence was excited at 275 nm and monitored at 305 nm. In order to follow the $Ca^{2+}$ dependent hydrophobic pocket opening in the N-domain of CaM, bis-ANS fluorescence was excited at 390 nm and monitored at 495 nm. To determine the $Ca^{2+}$ sensitivity of wtCAM or its mutants in the presence of human

RYR2 CaM-binding domain, intrinsic Trp fluorescence of RyR2 peptide was excited at 295 nm and monitored at 330 nm as microliter amounts of $CaCl_2$ were added to 2 ml of RyR2 peptide (1.5 μM) in the presence of 0.5 μM of wtCaM or its mutants in the titration buffer. Free [$Ca^{2+}$] was calculated using a program, EGCA02, developed by Robertson and Potter[66]. Each reported $pCa_{50}$ represents an average of at least three titrations ± S.E. To determine the apparent affinity of wtCaM or its mutants for RYR2 CaM-binding domain, microliter amounts of wtCaM or its mutants were added to 2 mL of titration buffer containing 0.5 μM of RyR2 peptide in either saturating (pCa=3.85) or sub-saturating $Ca^{2+}$ (pCa=6.6). Each reported $pCaM_{50}$ represents an average of at least three titrations ± S.E. The data were fit with a logistic sigmoid function, mathematically equivalent to the Hill equation, as previously described[82-84].

***Kinetic Fluorescence Measurements.***

All kinetic measurements were carried out using an Applied Photophysics Ltd. (Leatherhead, UK) model SX.18MV stopped-flow apparatus with a dead time of ~1.4 ms at 20 °C. The samples were excited using a 150W xenon arc source. The stopped-flow buffer for all experiments consisted of 10 mM MOPS, 150 mM KCl (pH 7.0). Tyr fluorescence was excited at 275 nm with emission monitored through a UV-transmitting black glass (UG1) filter from Oriel (Stratford, CT, USA). Trp fluorescence was excited at 295 nm with emission also monitored through a UG1 filter. Quin2 fluorescence was exited at 330 nm with emission monitored through a 510 nm broad band pass interference filter from Oriel (Stratford, CT, USA). Bis-ANS fluorescence was excited at 390 nm with emission monitored through a 500 nm long pass interference filter from Newport (Irvine, CA, USA).

In order to measure rates of $Ca^{2+}$ dissocation from the C-domain of wtCaM or its mutants via conformational change, 3 μM CaM in stopped-flow buffer plus 200 μM $Ca^{2+}$ was rapidly mixed with 10 mM EGTA in stopped-flow buffer. To measure the $Ca^{2+}$ dependent rate of the N-domain hydrophobic pocket closure, 1 μM CaM and 1 μM Bis-ANS in stopped-flow buffer plus 200 μM $Ca^{2+}$ was rapidly mixed with 10 mM EGTA in stopped-flow buffer. The fluorescent $Ca^{2+}$ chelator Quin2 was utilized to determine the rate of $Ca^{2+}$ dissociation from wtCaM or its mutants in the absence or presence of the RyR2 peptide. Each CaM protein (3-6 μM) in the absence or presence of RyR2 peptide (3-fold excess over CaM) in the stopped-flow buffer plus 30 μM $Ca^{2+}$ was rapidly mixed with an equal volume of Quin-2 (150 μM) in the stopped-flow buffer. The data were fit using a program by P. J. King, Applied Photophysics Ltd. that utilizes the nonlinear Levenberg-Marquardt algorithm. Each $k_{off}$ represents an average of at least three separate experiments ± S.E., each averaging at least three shots, which were fit with a double exponential equation since Quin2 reports the rates of $Ca^{2+}$ dissociation from both the N- and the C-domains of CaM.

***Intact RyR2 Affinity***

To resolve CaM binding affinity for intact RyR2 in SR membranes, we used a FRET–based competition binding assay, as previously[85] (Hwang 2014). Briefly, SR vesicles from porcine ventricular myocardium[86] (Fruen 2000) were incubated with 70 nM AlexaFluor488-labeled FKBP12.6 to attach a fluorescent donor probe on RyR2. Unbound donor-FKBP was removed as supernatant following SR sedimentation via centrigation. Assay conditions included 3 mg/mL of Donor-FKBP-SR, 0.001 – 10 μM WT or mutant CaM, 50 nM AlexaFluor568-labeled CaM, 20 mM K-PIPES (pH 7.0), 150 mM KCl, 5 mM reduced glutathione, 1 mM EGTA, 0.1 mg/mL BSA, 1 μg/mL aprotinin/leupeptin, and 1.02 mM CaCl2 to yield 30 μM free [$Ca^{2+}$]. After a 2.5-hour incubation at 25°C, the samples were transferred to a 384-well plate and fluorescence lifetime was acquired using a Fluorescence Innovations Inc fluorescence lifetime plate reader with 473nm Excitation and 520/17nm emission filter[87,88] (Petersen 2014, Rebbeck 2017). FRET was calculated based on the fractional decrease of donor fluorescence lifetime ($\tau_D$) in the presence of acceptor ($\tau_{DA}$), according to FRET=1–$\tau_{DA}/\tau_D$.

***Myocyte Isolation***

In this study, we used adult (3-6 months old), male wild type (WT) and genetically modified C57BL/6 mice, $RyR2^{S2814D +/+}$ [89]. Animal use was approved by the Animal Care and Use Committee of Vanderbilt University, USA (animal protocol #M1600259-00), the Ohio State University (animal protocol 2010A00000117-R5), and Mississippi State University (animal protocol #25-466) in accordance with NIH guidelines. Mouse ventricular myocytes were isolated as previously described[18], after mice were fully anesthetized using 4% isoflurane in 95% oxygen. The hearts were quickly excised and perfused on a Langendorff's apparatus at 37°C for 5 minutes of perfusion with nominally $Ca^{2+}$-free Tyrode solution (containing, in mM: 140 NaCl, 5.4

KCl, 0.5 $MgCl_2$, 10 HEPES, and 5.6 glucose [pH 7.3]). They were then digested with Liberase TH containing Tyrode solution. Cells were washed twice by gravity sedimentation for 20 minutes in standard Tyrode solution containing 0.2 mM $CaCl_2$. The final suspension contained 0.6 mM $Ca^{2+}$. Cells were plated onto laminin-coated glass cover slips and allowed to attach for 30 minutes at room temperature.

***$Ca^{2+}$ Imaging in Permeabilized Cardiomyocytes***

Attached cells were permeabilized with saponin (40 µg/mL) for 45 seconds and then bathed for 30 minutes in a freshly-made internal solution (pH = 7.2) containing: K-aspartate (120 mM), KCl (15 mM), K2HPO4 (5 mM), $MgCl_2$ (5.6 mM), HEPES (10 mM), dextran (4% w/v), MgATP (5 mM), phosphocreatine-Na2 (10 mM), creatine phosphokinase (10 U/mL), reduced L-glutathione (10 mM), $CaCl_2$ (0.12 mM), CaM autocamtide-2-related inhibitory peptide (AIP, 1 µM), and Fluo-4 (0.03 mM). Additionally, the internal solution contained 0.5 or 0.1 mM EGTA (free [$Ca^{2+}$] ~50 nM or ~120 nm, respectively) for recording of $Ca^{2+}$ sparks or waves, respectively. In experiments utilizing recombinant CaM proteins, permeabilized myocytes were incubated with either wtCaM or its mutants for 25 minutes to allow equilibration of CaM binding to its targets prior to recording of $Ca^{2+}$ sparks and waves. Fluo-4 was excited with the 488 nm line of an argon laser and emission was collected at 500 to 600 nm. Fluo-4 fluorescence was recorded in the line-scan mode of the confocal microscope (Zeiss LSM 510, Olympus Fluoview 1000, or Nikon Ti2 AXR confocal). Spark analysis was performed in ImageJ with the SparkMaster plugin using a background setting of 5 and criteria of 3.8. Spark mass was calculated from the equation: spark mass = 1.206 x Amplitude x $FWHM^3$. The 1.206 value is a geometrical conversion factor that can be used for converting the Full Width Half Max (FWHM) distance parameter into a volume. Mass = $A \times [\Pi/(4 \times \ln 2)]^{3/2} \times FWHM^3$ and therefor volume can be approximated by 1.206 x Amplitude x FWHM. For a detailed description of the derivation please see [9073]. Leak was calculated from the equation: leak = frequency x mass. Statistical analysis was performed using Prism v10 (GraphPad Software, Inc.). The 4 categories of spark data (frequency, amplitude, mass and leak) were analyzed using a one tailed Mann -Whitney test that does not assume a gaussian distribution. A p-value of 0.05 was used as the threshold to reject the null hypothesis.

#### 7. SUPPLEMENTARY FIGURES

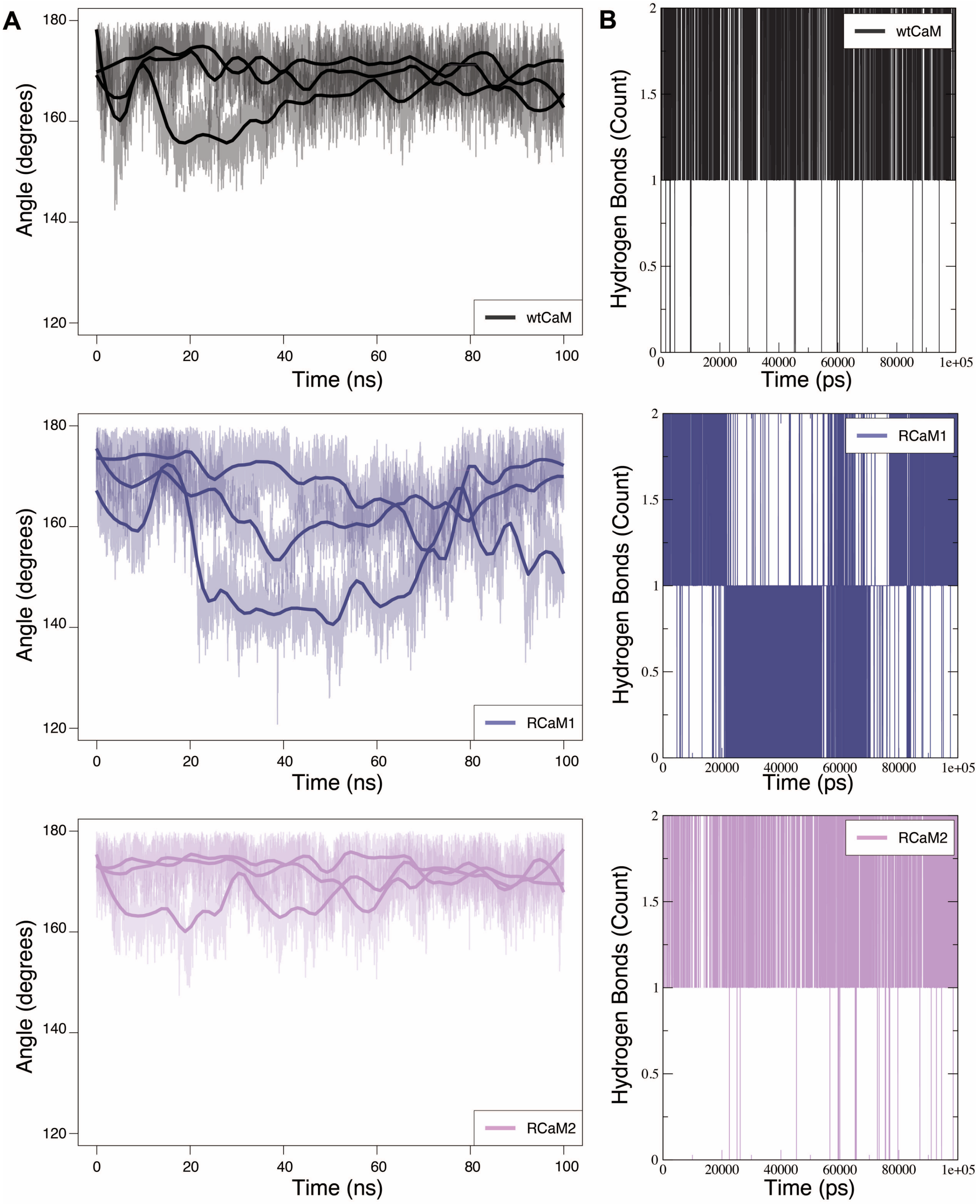


**Supplementary Figure S1. Analysis of RyR2 CaM-binding peptide geometry and intramolecular hydrogen bonding.**

(A) Time-dependent variation in the angle formed by Cα atoms of RyR2 residues 3589, 3592, and 3595 over 100-ns molecular-dynamics simulations of wtCaM (black), RCaM1 (blue), and RCaM2 (magenta) complexes. Thick lines represent moving averages; shaded regions denote instantaneous fluctuations. (B) Time evolution of intramolecular hydrogen bonds within the α-helical segment of the RyR2 peptide (residues 3590–3595) during a representative trajectory. Each vertical line indicates the presence of one or two hydrogen bonds at a given time point.

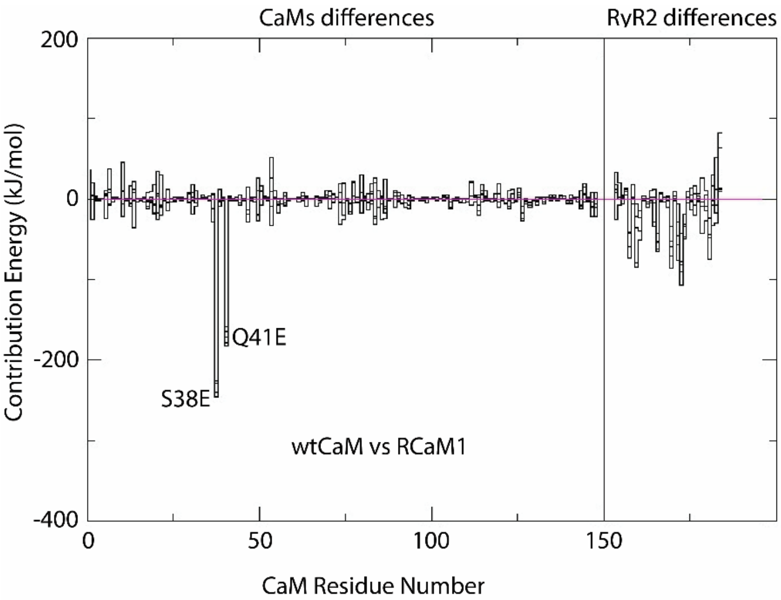


**Supplementary Figure S2. Differences in average per-residue contributions to the Δ binding energy between wtCaM and RCaM1.** The plot shows the difference in average binding energy contribution of each residue in CaM and RyR2 between wtCaM–RyR2 and RCaM1–RyR2 complexes, calculated across all simulation trajectories at 1-ns intervals. Residues S38E and Q41E correspond to redesigned positions in the RCaM1 variant.

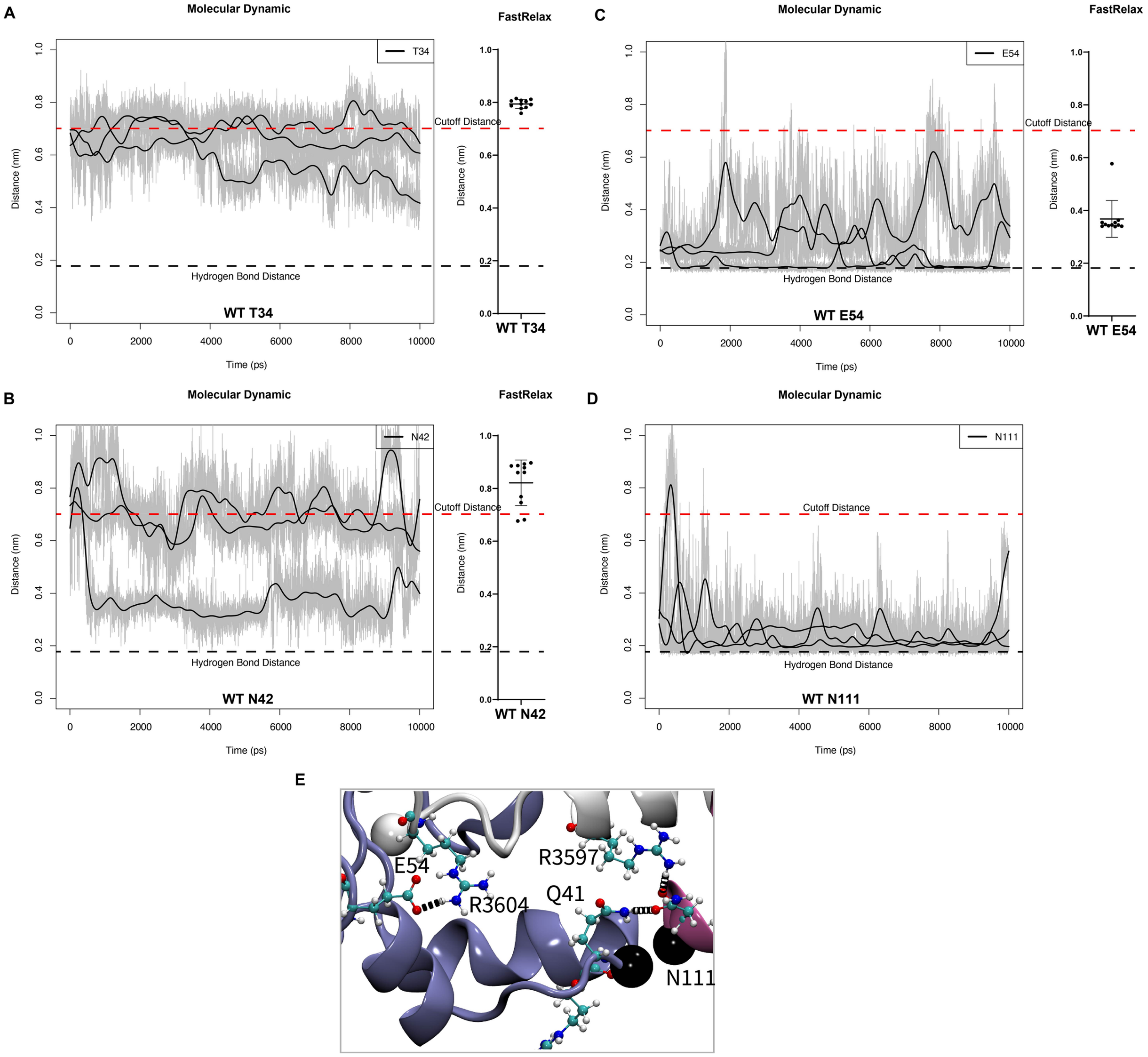


**Supplementary Figure S3.**
(A-D) Time-dependent changes in the distance between selected wtCaM residues and the nearest atom of the RyR2 peptide during molecular dynamics simulations (left). Distances are compared with those observed in the corresponding crystal structure and in the eight top-ranked FastRelax models (right). The dashed red line indicates the 7 Å cutoff distance used for selecting residues for redesign; the dashed black line marks the typical hydrogen bond distance.
(E) Structural snapshot illustrating spatial connections between residues N111, Q41, and R3597 within the annealed wtCaM–RyR2 complex.

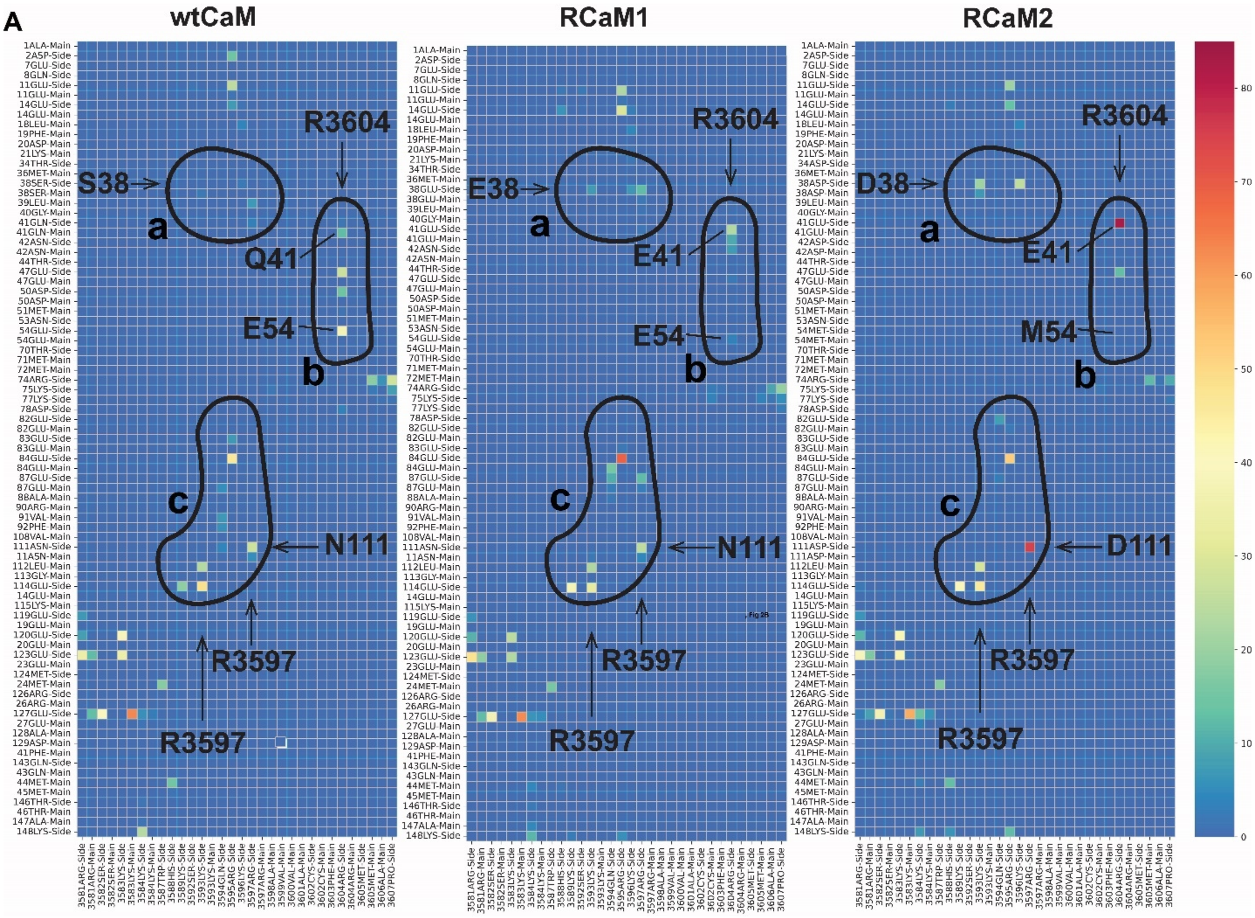


**Figure S4. Bonding Network Map.**
The heatmaps show the percentage of molecular dynamics simulation time during which specific hydrogen or salt-bridge interactions were maintained. Red indicates interactions that persisted for a larger fraction of the trajectory, whereas blue denotes the absence of bonding. Network regions involving redesigned residues are outlined and labeled “a,” “b,” and “c.” Arrows mark key residues (S38/D38, Q41/E41, E54/M54, and N111/D111) and their RyR2 interaction partners (R3604, R3597).

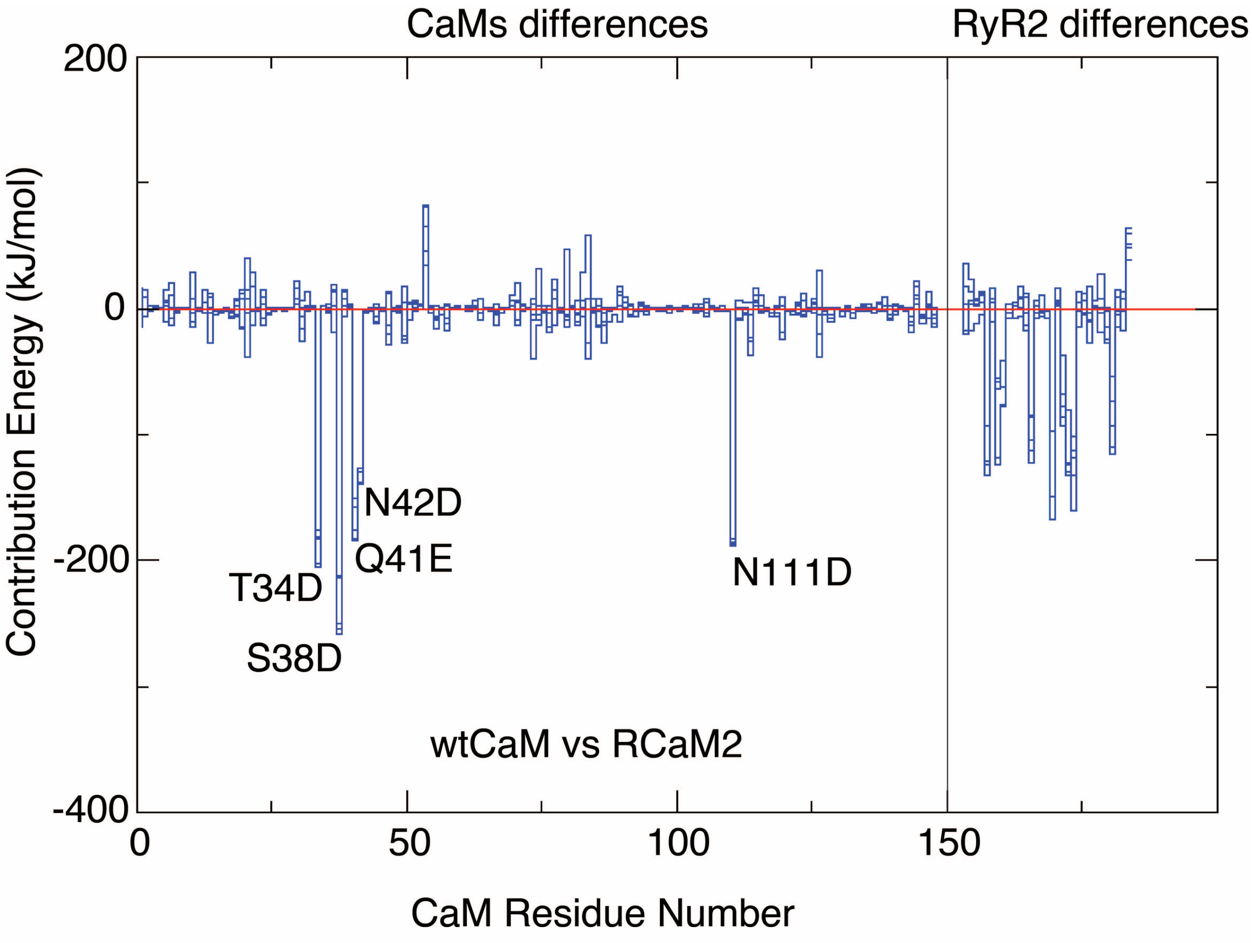


**Supplementary Figure S5. Differences in average per-residue contributions to the Δ binding energy between wtCaM and RCaM2.**
The plot shows differences in average per-residue binding-energy contributions for CaM and RyR2 between wtCaM–RyR2 and RCaM2–RyR2 complexes, calculated over all molecular-dynamics trajectories. Residues T34D, S38D, Q41E, N42D, and N111D correspond to redesigned positions in the RCaM2 variant and represent the main contributors to altered binding energetics.

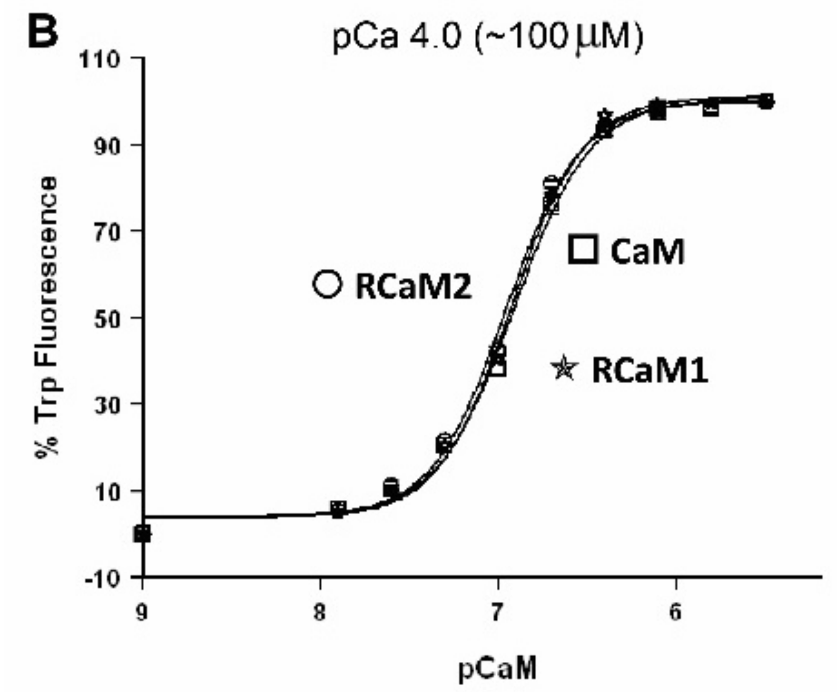


**Supplementary Figure S6. CaM affinity for the RyR2 peptide under $Ca^{2+}$-saturated conditions.** The plot shows Trp fluorescence measurements used to assess the apparent binding affinity of wtCaM, RCaM1, and RCaM2 for the RyR2 CaM-binding peptide in the presence of saturating $Ca^{2+}$ (pCa 4.0, ~100 µM). The fluorescence signal reflects the concentration of CaM required for 50% peptide binding.

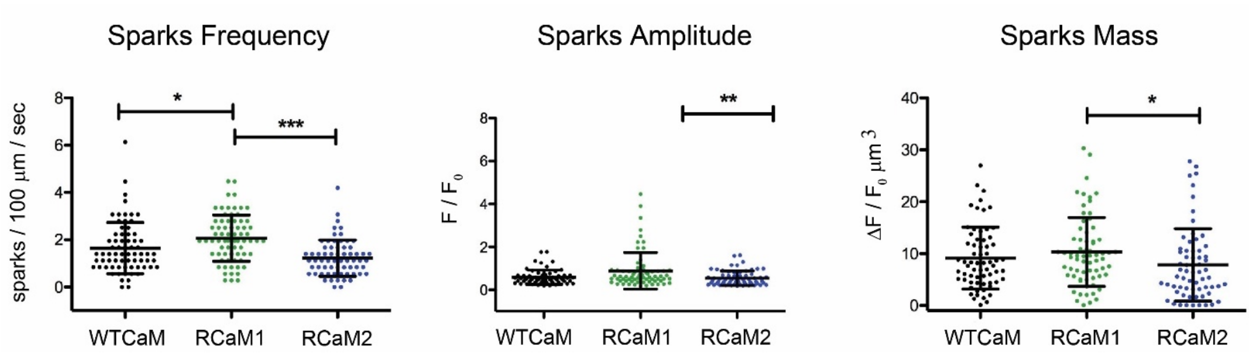


**Supplementary Figure S7. $Ca^{2+}$ Spark Properties in S2814D Ventricular Myocytes Incubated with wtCaM, RCaM1, or RCaM2.**

Calcium sparks were recorded in saponin-permeabilized ventricular cardiomyocytes from S2814D mice using confocal microscopy and Fluo-4 fluorescence. Cells were incubated with 1 μM wtCaM, RCaM1, or RCaM2 30 minutes prior to imaging. (Left) Spark frequency shows increased activity with RCaM1 and reduced activity with RCaM2 relative to wtCaM. (Middle) Spark amplitude exhibits a modest increase with RCaM1, with values for RCaM2 comparable to wtCaM. (Right) Spark mass parallels these differences. Each point represents an individual spark event; data are shown as mean ± SD.

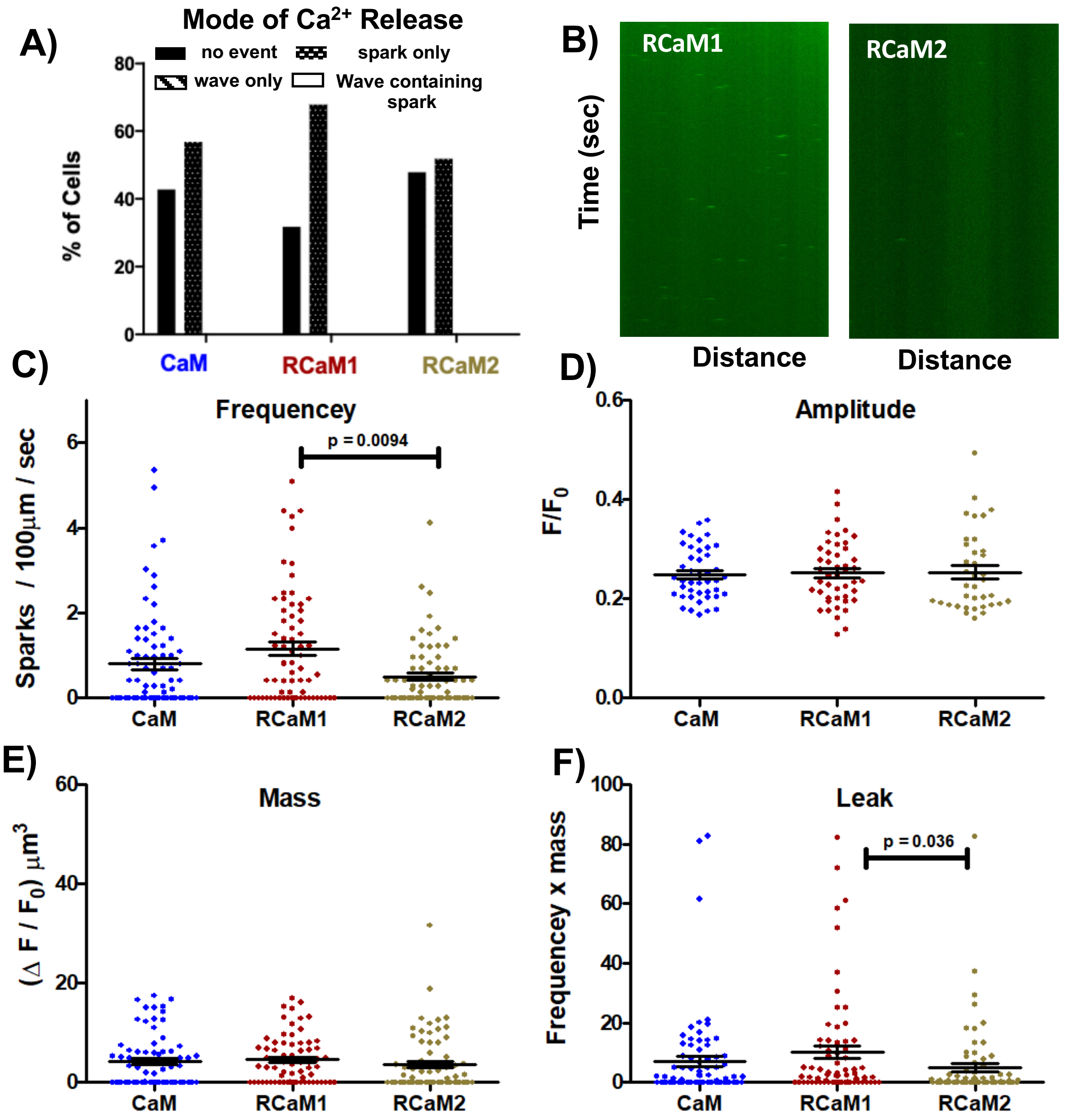


**Supplementary Figure S8. Ex vivo $Ca^{2+}$ spark properties in permeabilized ventricular myocytes expressing wild-type RyR2 at 50 nM free $Ca^{2+}$.**

(A) Modes of $Ca^{2+}$ release across cells incubated with wtCaM, RCaM1, or RCaM2.

(B) Representative line-scan images illustrating increased spark activity with RCaM1 and reduced activity with RCaM2.

(C) Spark frequency (sparks per 100 μm per second) showing increased activity with RCaM1 and decreased activity with RCaM2 ($p < 0.01$).

(D–E) Spark amplitude and mass, respectively, showing no significant differences across conditions.

(F) Spark leak (frequency × mass) demonstrating parallel differences consistent with frequency changes.

#### 4. SUPPLEMENTARY MOVIE LEGEND

*Supplementary movie SM1 provided as a separate file in mp4 format*

**Supplementary Movie SM1. MD simulation showing the conformational behavior of the $Ca^{2+}$-bound RCaM1 (blue N-lobe, pink C-lobe) bound to the RyR2 CaM-binding peptide (gray).** Throughout the trajectory, RCaM1 samples an "unlocked" configuration in which the N- and C-lobes separate, inducing bending of the RyR2 peptide near Ser3592 (not shown). This peptide deformation corresponds to the closed-ready, leak-prone state observed in leaky RyR2 variants. The movie highlights the dynamic contrast between RCaM1 and the compact, peptide-stabilizing "annealed" conformations favored by wtCaM and RCaM2 (not shown). Magenta spheres indicate C$\alpha$ atoms of residues 40 and 113 in RCaM1. Solvent and $Ca^{2+}$ ions are omitted for clarity.
